# Vasculature-associated adipose tissue macrophages dynamically adapt to inflammatory and metabolic challenges

**DOI:** 10.1101/336032

**Authors:** Hernandez Moura Silva, André Báfica, Gabriela Flavia Rodrigues-Luiz, Jingyi Chi, Patricia d’Emery Alves Santos, Bernardo S. Reis, David P. Hoytema van Konijnenburg, Audrey Crane, Raquel Duque Nascimento Arifa, Patricia Martin, Daniel Augusto G.B. Mendes, Daniel Santos Mansur, Victor Torres, Ken Cadwell, Paul Cohen, Daniel Mucida, Juan José Lafaille

**Affiliations:** Kimmel Center for Biology and Medicine at the Skirball Institute; New York University School of Medicine, New York, NY, USA; Laboratório de Imunobiologia, Departamento de Microbiologia, Imunologia e Parasitologia, Centro de Ciências Biológicas, Universidade Federal de Santa Catarina, Santa Catarina CEP 88040-900, Brazil; Laboratory of Mucosal Immunology, The Rockefeller University, New York, NY, USA; Laboratory of Molecular Metabolism, The Rockefeller University, New York, NY, USA; Department of Microbiology, New York University School of Medicine, New York, NY, USA; Department of Pathology, New York University School of Medicine, New York, NY, USA

**Author notes:** Corresponding Authors (H.M.S); (J.J.L.). Correspondence address: 430 E 29th street. 4th floor, Lab 460A. New York, NY. USA. 10016.

## Abstract

Tissue-resident macrophages comprise the most abundant immune cell population in healthy adipose tissue. Adipose tissue macrophage populations change during metabolic stress and ageing, and are thought to contribute to the pathogenesis of obesity. Here, we studied adipose tissue macrophage subpopulations in the steady state, and in response to nutritional and infectious challenges.

Using comprehensive cell-surface-based and gene expression analyses, we found that tissue-resident macrophages from healthy epididymal white adipose tissue (eWAT) tightly associate with blood vessels, displaying a very high endocytic capacity. We refer to these cells as Vasculature-associated Adipose tissue Macrophages (VAMs). Chronic high fat diet (HFD) feeding results in the accumulation of a monocyte-derived CD11c^+^CD64^+^ double positive (DP) macrophage eWAT population with a predominant anti-inflammatory gene profile, but reduced endocytic function. In contrast, fasting rapidly and reversibly leads to VAM depletion, while acute inflammatory stress induced by pathogens transiently depletes VAMs and simultaneously boosts DP macrophage accumulation. Our results indicate that adipose tissue macrophage populations adapt to metabolic stress and inflammation, suggesting an important role for these cells in restoring homeostasis.

## Introduction

The adipose tissue of healthy animals contains a large population of innate and adaptive immune cells, numerically dominated by macrophages (Lumeng et al., 2007; Weisberg et al., 2003; Xu et al., 2003). The functions of these cells in the healthy adipose tissue of animals in steady state are not completely understood, given that there is no apparent insult. However, the quantity, variety and distribution of these immune cells suggest a critical role in maintaining homeostasis of the tissue. The proportions and numbers of these immune cell types is known to change extensively in conditions such as high fat diet (HFD), genetically-determined obesity and aging, among other situations.

It has been long known that obesity is associated with a heightened inflammatory condition, in which TNFα is an important mediator of insulin-resistance (Feinstein et al., 1993; Hotamisligil et al., 1995; Hotamisligil et al., 1993). It is thought that the myeloid cells found in the adipose tissue during diet-induced obesity (DIO) contribute to the pathogenesis of insulin resistance and the metabolic syndrome, by increasing their secretion of pro-inflammatory cytokines, such as TNFα. However, it is unclear how adipose immune cells mediate pathology, and what the possible roles of myeloid cells are in the steady state (Dalmas et al., 2011; Fitzgibbons and Czech, 2016; Lumeng, 2013; Wynn et al., 2013; Xu et al., 2013).

HFD induces obesity and insulin resistance in B6 mice (Rebuffe-Scrive et al., 1993; Surwit et al., 1988). Adipocytes become hypertrophic, which could be the initial signal to trigger inflammation (Rutkowski et al., 2015; Sun et al., 2011); however, cultured adipocytes made hypertrophic with oleic acid, display insulin resistance without triggering an inflammatory response (Kim et al., 2015). Thus, the quantity and quality of the inflammatory response in obesity-related disease is not clearly established. Moreover, wild animals endure fluctuations in food supply, in which the adipose tissue triglyceride reserves are mobilized via lipolysis or built from the diet, and little is known about the response of immune cells to this natural instability.

In this report, we found that tissue-resident macrophages from healthy epididymal white adipose tissue (eWAT), a visceral fat depot, not only surround adipocytes but also associate tightly with blood vessels, being among the most endocytic cells in the body for blood-borne molecules. We refer to these macrophages as Vasculature-associated Adipose tissue Macrophages (VAMs). Upon chronic high fat diet (HFD) feeding, a monocyte-derived population of macrophages, poorly represented in animals fed a normal diet (ND), surges in the eWAT. Contrary to expectations, these macrophages have predominantly an anti-inflammatory and tissue repair/healing gene signature, with no expression of typical M1 genes. Importantly, eWAT macrophages from HFD-fed mice have a reduced endocytic capacity. Acute inflammatory stress, such as that induced by LPS administration or *Salmonella* infection, results in a rapid and transient reduction in the main populations of VAMs, with an increase in monocyte-derived macrophages, whereas a brief fasting period rapidly and reversibly reduces all macrophage populations, especially VAMs. Thus, the macrophage populations adapt to metabolic stress and inflammation by adjusting their numbers and gene expression, even when pathology ends up prevailing.

## Results

### White adipose tissue in steady state has multiple populations of macrophages

We characterized the epididymal white adipose tissue (eWAT) lymphoid and myeloid cell populations under steady state. We chose to focus on the visceral WAT because excess abdominal fat, unlike excess subcutaneous fat, strongly correlates with metabolic syndrome (Sharma, 2002; von Eyben et al., 2003). First, we carried out an extensive flow cytometry analysis in perfused male mice at age 12-20 weeks. Flow cytometry of adipose tissue macrophages is made somewhat difficult because of their high autofluorescence. To minimize the impact of autofluorescence, we carefully selected the fluorochromes for each cell marker, and used fluorescence minus one controls (FMOs). The gating strategy, including all markers, is shown in **Fig. S1A-L**, and the abbreviated list of markers used throughout this manuscript to identify the key populations is shown in **Fig. 1A**. A useful series of surface markers for the eWAT myeloid populations is shown in **Fig. S1M**. We observed a minor Ly6C^HIGH^ monocyte population (**Fig. S1K**), two populations of dendritic cells (DC), and four populations of macrophages, besides eosinophils and neutrophils (**Fig. S1J**), innate lymphoid cells (**Fig. S1D-E**), and lymphocytes (**Fig. S1F**). The dendritic cells belong to the two main tissue DC populations, CD11b^-^CD103^+^ and CD11b^+^CD103^-^ DC (**Fig. S1I, S1L and S1M**). As explained in further detail below, we refer to the macrophage populations as Vasculature-associated Adipose tissue Macrophages (abbreviated VAM) 1, VAM2, preVAMs, and double positive (DP) macrophages. VAM1 and 2 are very similar, but can be readily distinguished by their surface expression level of MHC Class II and Tim4 (**Fig. S1H**). The composition of all immune cells in the healthy male eWAT is summarized in **Fig. S2A**. The combination of the two VAM populations outnumbers the sum of all other CD45^+^ eWAT cells. CD11b^+^ DC and eosinophils are, respectively, the third and fourth cell types by abundance. DP macrophages, which are quite rare in eWAT from mice fed ND (**Fig. S2A**), expresses high surface levels of CD11b, CD11c, CD64, MHC II, and intermediate levels of CD206. We refer to these macrophages as DP because of their surface expression of CD11c and CD64. CD64 expression distinguishes DP macrophages from CD11b^+^ DC, and CD11c expression distinguishes DP macrophages from preVAM PreVAMs and DP macrophages express lower level of surface CD206 than VAMs (**Fig. S1 L-M**). Thus, the myeloid populations form the eWAT are more complex than previously anticipated.

**Figure 1:**
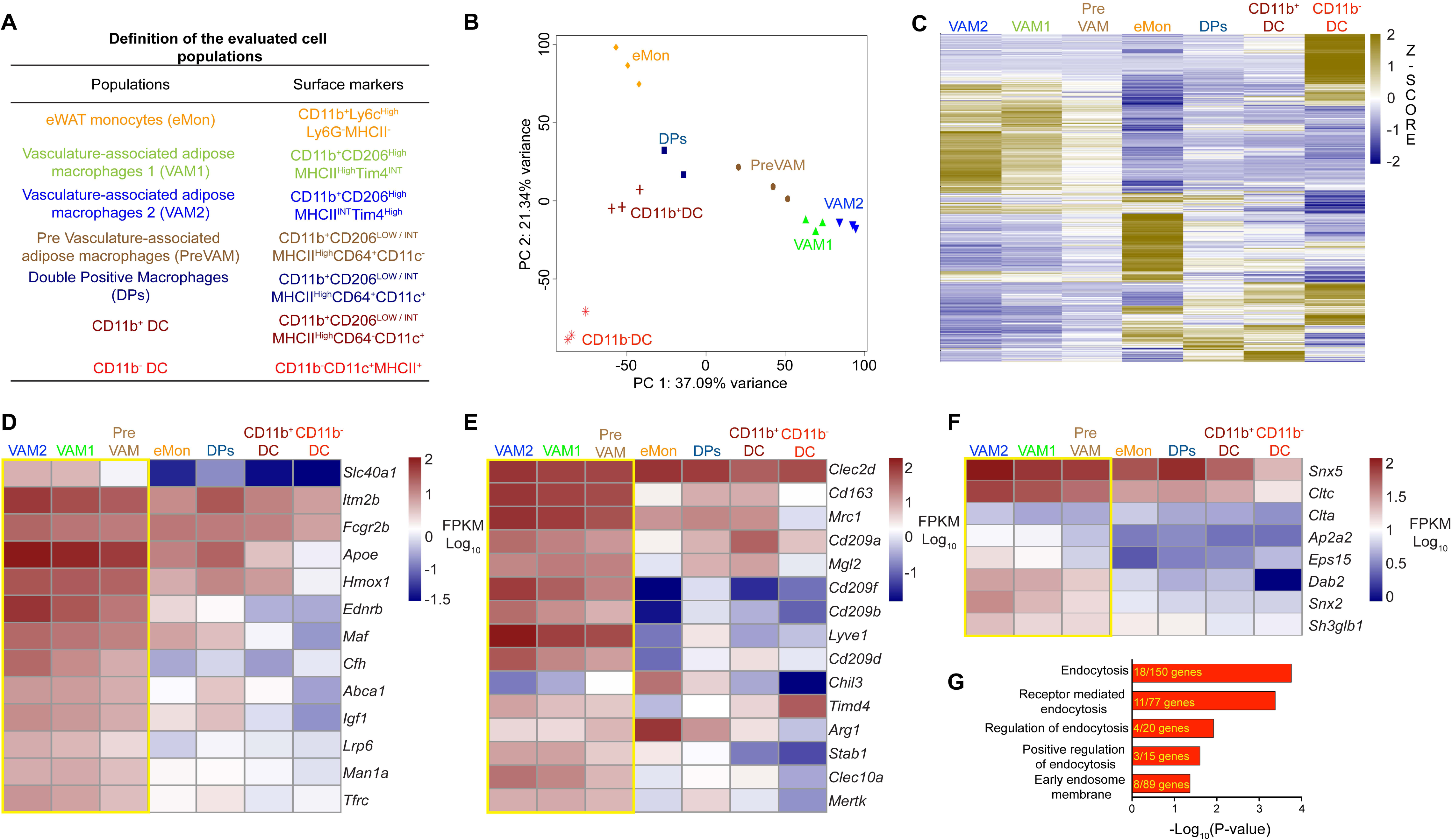
The predominant eWAT macrophage subpopulations display a gene signature enriched in anti-inflammatory, scavenging, and endocytosis-related pathways. **(A)** Abbreviated description of the key markers defining the cell populations analyzed. The colors identifying each population are used throughout the manuscript. **(B)** Principal component analysis of the transcriptional signature obtained by RNAseq data from the populations described in **A**. Adult (16 week-old) WT males were used. Each dot represents data from a pool of 2-3 epididymal fat pads. **(C)** Gene expression profile of eWAT myeloid subpopulations described in A, showing all genes for broad visualization. Each column represents consolidated data from 3 biological replicates per cell type. The z-score of the gene expression profiles gives a scale to measure the differential expression. **(D-F)** Heatmap of genes differentially expressed in VAMs highlighting: **(D)** their active, anti-inflammatory, insulin-sensitizing, repair and detoxifying gene signature; **(E)** genes encoding lectin and scavenger receptors; **(F)** genes associated with endocytosis. **(G)** VAM gene signature, analyzed by functional enrichment, reveals an association with endocytic activity. Gene expression in D-F is displayed (FPKM in the Log_10_ base). Yellow boxes mark the VAMs and preVAM populations.

Given the different pathophysiological roles of the various fat depots, we compared the composition of myeloid cells in eWAT, mesenteric visceral WAT (mWAT), and subcutaneous WAT (sWAT). VAM1 and VAM2 are significantly enriched in the eWAT compared to the other WAT depots, and the subcutaneous WAT had an appreciably lower number of total macrophages and dendritic cells than the two visceral depots (**Fig. S2B**).

To gain information regarding cell morphology, we sorted the populations and observed them in Giemsa-stained cytospin preparations. The most remarkable aspect is that, in healthy mice eating a normal diet (ND), VAMs and preVAMs have abundant lipid droplets in their cytoplasm (**Fig. S3**), a morphology that resembles foam cells in atherosclerotic vessels, and adipose tissue macrophages from obese mice (Aouadi et al., 2014), but, to our knowledge, not previously reported in adipose macrophages from lean mice. In ob/ob mice, reducing lipid droplet content via Lpl knockdown resulted in a worsening of glucose tolerance (Aouadi et al., 2014). This morphological feature of VAMs and preVAMs is consistent with an active protective role during steady-state in lean mice.

We used RNASeq to carry out a broad transcriptional analysis of eWAT monocytes, macrophages and dendritic cells. Principal component analysis showed that VAM1, VAM2, and preVAMs cluster near each other (**Fig. 1B**). CD11b^-^ DC and eWAT monocytes (eMon) are distant from the other myeloid cells analyzed. A global heatmap shows the similarities and differences between the eWAT cell types studied (**Fig. 1C**). **Table S1** contains the list of all genes discussed in this manuscript, including the official names, aliases and synonyms.

The small population of adipose monocytes (eMon) has many distinguishing gene expression features compared to Ly6C^+^ and Ly6C^-^ circulating monocytes (Mildner et al., 2017), suggesting a rapid adjustment to the adipose tissue environment (**Fig S4A**). They seem to be poised to differentiate into macrophages in the eWAT, as indicated by their high expression *Mafb*, which has been shown to be essential in the differentiation of monocyte-derived macrophages (Goudot et al., 2017). They are mostly cells in transition, although displaying typical monocyte cell surface markers. Some of the genes expressed at low level by blood monocytes that are highly expressed by eWAT monocytes are *Dab2, Nfkbiz, Hivep2, Skil, Egr1, Rel, Plek, Rbpj, Odc1*, and *Clec7a* (**Fig. S4B**). These genes are also expressed by eWAT macrophages and DC, suggesting the adaptation of monocytes to the eWAT environment. In contrast, some genes expressed at high level by blood monocytes are either not expressed, or expressed at very low levels by eWAT monocytes, indicating that they are rapidly repressed. Some examples include: *Zfp414, Hmg20b, Drap1, Tsc22d4*, and *Bmyc* (**Fig. S4B**).

Gene expression confirmed that the eWAT contains the two typical populations of dendritic cells (DC), a CD103^+^ (*Itgae*) CD11b^-^ population, and a CD103^-^CD11b^+^ population. CD11b^-^ DC express high levels of the genes *Xcr1, Irf8, Id2, Mycl, Notch2, Tlr3, Clec9a, Ly75* and *Zbtb46*, and a low level of *Irf4*. CD11b^+^ DC express relatively high level of *Irf4* (highest among eWAT monocytes, macrophages and DC) and a low level of *Irf8*, with moderate *Notch2* mRNA expression. *Zbtb46* expression levels in CD11b^+^ DC are intermediate between the CD11b^-^ DC and the macrophages. This CD11b^+^ DC population also displays low surface F4/80, which is higher than CD11b^-^ DC but lower than the four macrophage populations (**Fig. S1M**). CX3CR1 expression is mixed in CD11b^+^ DC (**Fig. S1M**), perhaps reflecting the known heterogeneity of this population (Mildner and Jung, 2014). Both eWAT DC populations express similarly high levels of the *Clec12a* gene, and neither expresses *Esam*, two markers that have been used to distinguish among DC populations in the spleen and intestine (Lewis et al., 2011) (**Fig. S4C)**.

After this initial characterization of all the myeloid cells populations in the eWAT, we focused most of our subsequent studies on the macrophage populations. It has been established that these are tissue-resident macrophages that proliferate locally, and are independent of recently recruited monocytes (Amano et al., 2014). In fact, total genetic ablation of the gene encoding MCP-1 (*Ccl2*), the key chemokine involved in the recruitment of monocytes, did not reduce the number of adipose tissue macrophages (Inouye et al., 2007; Kirk et al., 2008).

Our gene expression analysis suggests that steady-state VAM1 and VAM2 macrophages in mice eating a normal diet (ND) are very active, and have anti-inflammatory, insulin-sensitizing, repair and detoxifying functions (**Fig. 1D**). Confirming and expanding the flow cytometry data, they express high levels of genes encoding the lectin and scavenger receptors, and also genes encoding “eat me” signals for apoptotic cells (**Fig. 1E**)

VAMs 1 and 2 appear to be well equipped for endocytosis in steady state conditions. They express high levels of the genes encoding Clathrin heavy chain (*Cltc*), the Clathrin alpha light chain (*Clta*) genes, the clathrin adaptor protein *Dab2*, and components of the clathrin-associated Ap2 complex, such as *Ap2a2* and *Ap2b1*. Another endocytosis adaptor highly expressed by VAMs was *Eps15*, which has been implicated both in clathrin-dependent and independent endocytosis (Carbone et al., 1997; Sigismund et al., 2005) (**Fig. 1F**). Interestingly, VAMs also express high levels of Endophilin B1 (*Sh3glb1*), involved in ultrafast, clathrin-independent endocytosis, a mechanism utilized by very active cells (Watanabe and Boucrot, 2017). Genes involved in macrophage macropinocytosis, such as *Snx5* (Lim et al., 2015), are also very highly expressed. To function in the recycling of membrane proteins, SNX5 forms a heterodimer with SNX2 (Kvainickas et al., 2017; Simonetti et al., 2017), which is also highly expressed by VAMs (**Fig. 1F**).

Functional enrichment analysis (GO term analysis) strongly suggest that steady state VAMs have an active endocytic and filtering function (**Fig. 1G**). Taken together, these data suggest that VAMs are an abundant and active myeloid population in the eWAT, with a genetic signature strongly associated with a high endocytic capacity.

### VAMs in eWAT are intimately connected with blood vessels, and highly endocytic

Given the Gene Ontology indications, we decided to test experimentally the hypothesis that VAMs may be particularly apt at endocytosis of blood-carried substances. First, we observed by confocal microscopy that VAMs maintain close contact with blood vessels, hugging them tightly at some points (**Fig 2A**).

**Figure 2:**
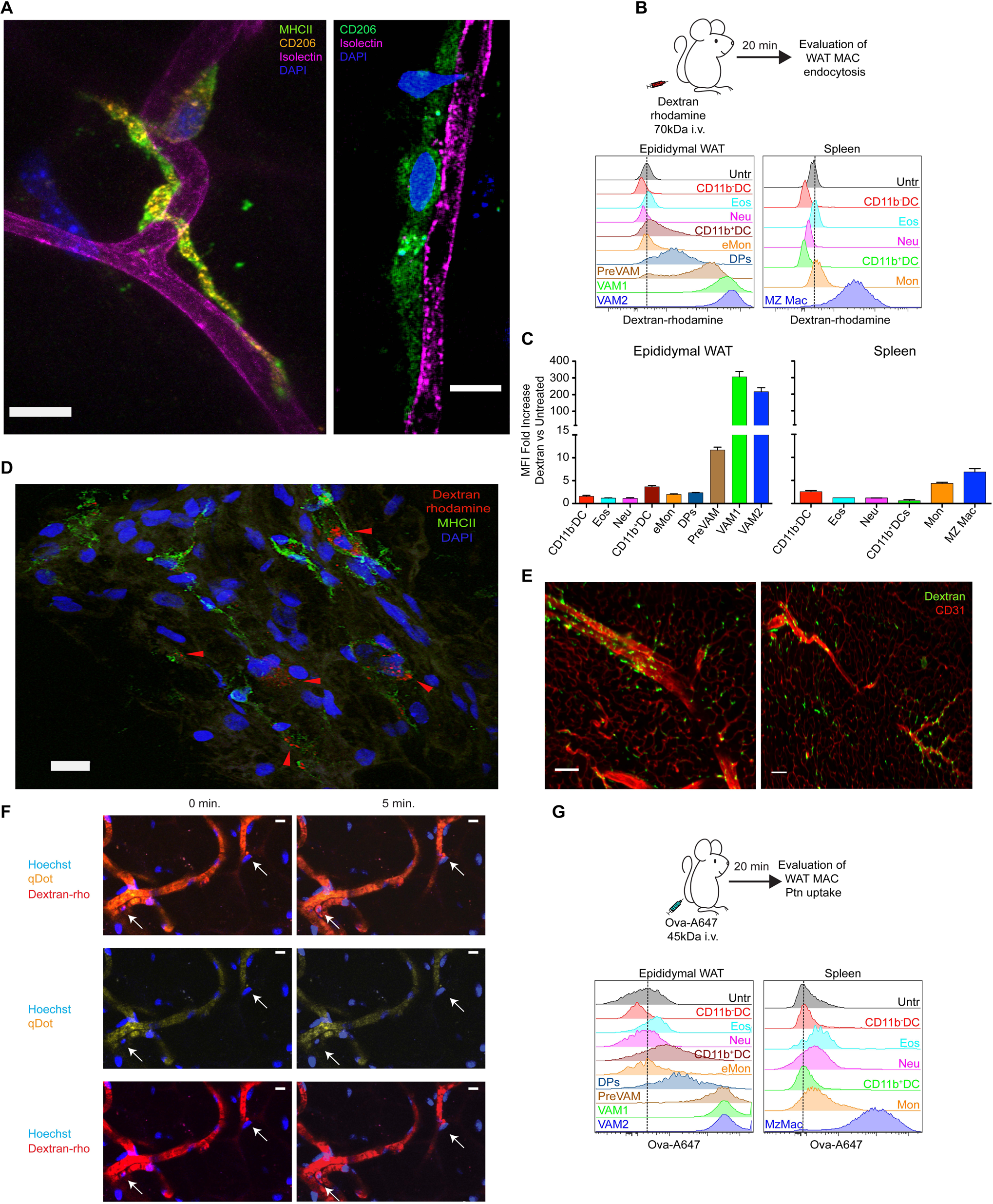
VAMs rapidly and efficiently uptake blood contents. **(A)** VAMs are intimately associated with WAT vessels. Representative confocal images of epididymal full mount sections showing the distribution of VAMs (CD206^+^MHCII^+^DAPI^+^) in close contact to blood vessels labeled with Isolectin. Scale bar =10 μm. 60x magnification. **(B)** Evaluation of VAMs endocytic activity *in vivo*. Dextran-rhodamine 70 kDa was injected i.v. Twenty min. post injection, animals were anesthetized and perfused intracardially. Endocytic activity was evaluated by measuring the increase of rhodamine fluorescence by flow cytometry. Histograms display the fluorescence intensity for rhodamine of eWAT and Splenic populations. Untr: Untreated. CD11b^-^DC (CD90^-^ CD19^-^CD11b^-^MHCII^+^CD11c^+^) Eos: Eosinophils (CD90^-^CD19^-^CD11b^+^SiglecF^+^). Neu: Neutrophils (CD90^-^CD19^-^CD11b^+^Ly6G^+^). CD11b^+^DC (CD90^-^CD19^-^CD11b^+^MHCII^+^CD11c^+^) Mon: Monocytes (CD90^-^CD19^-^CD11b^+^SiglecF^-^Ly6G^-^MHCII^-^Ly6C^HIGH^). MZ Mac: Marginal zone macrophages (CD90^-^ CD19^-^CD11b^INT^CD64^+^CD206^+^MHCII^+^). **(C)** Fold increase of the median rhodamine fluorescence intensity after injection (*n*=4)**. (D)** Representative confocal images of epididymal full mount sections showing that dextran-rhodamine is internalized in vesicles (arrowheads). Scale bar =10 μm. 40x magnification. **(E)** Light sheet microscopy of full epididymal fat pad post WAT clarification using Adipo-Clear technique (Chi et al., 2018) (**See Movie 1**). CD31 indicates blood vessels. Animals were injected with dextran-rhodamine, and after one hour submitted to sample preparation for Adipo-Clear. Both panels show VAMs around blood vessels from different z stacks. Scale bar = 1000 μm. 4x magnification. **(F)** Intravital microscopy analysis of eWAT dextran-rhodamine uptake five minutes after i.v. injection (**see Movie 2**). qDots (Qtracker 655) and Hoechst were i.v. injected immediately before the beginning of the recording, while dextran-rhodamine was injected while the recording was taking place. 25x magnification. Scale bar = 10 μm. **(G)** VAMs high endocytic capacity *in vivo* extends to proteins. Whole ovalbumin-A647 (Ova) was injected i.v. Twenty min post injection animals were analyzed as described in the legend of panel **B**. Ptn: Protein. MAC: Macrophage

To test endocytosis, we injected dextran-rhodamine 70 kDa intravenously, and, only 20 minutes after the i.v. injection, we sacrificed the mice and studied the myeloid populations by flow cytometry. VAM1 and VAM2 took up dextran very efficiently, followed by preVAMs. None of the two eWAT DC populations, or the eosinophils, significantly associated with Dextran sulfate under these conditions (**Fig. 2B and 2C**). Strikingly, eWAT VAMs took up dextran-rhodamine at a higher rate than spleen marginal zone macrophages (**Fig. 2B and 2C**). Note that marginal zone macrophages are directly exposed to the splenic blood sinuses and are specialized in the capture of blood-borne substances and, in particular, glycans, and express many of the same scavenger receptors that eWAT VAMs do (Mebius and Kraal, 2005). This indicates a very special property of eWAT VAMs. To ensure that the dextran-rhodamine was really internalized and not simply surface-associated, we carried out a confocal analysis, which shows that, indeed, dextran-rhodamine is within VAM vesicles (**Fig. 2D**). We did not observe any dextran-rhodamine uptake by CD45-negative cells (not shown).

Adipose tissue is typically difficult to image in 3D, because of high autofluorescence levels. To study whether the distribution of macrophages and blood vessels was uniform throughout the entire fat pad, we used a recently published whole adipose 3D immunolabeling and clearing method called AdipoClear (Chi et al., 2018). In order to label VAMs, we injected dextran-rhodamine i.v., and, one hour later, we perfused and fixed the animal, and dissected the epididymal fat pad for tissue clearing and whole-mount imaging with a light sheet fluorescence microscope. VAMs displayed a strong perivascular localization, which was uniform throughout the fat pad (**Movie 1 and Fig. 2E**), indicating that the entire fat pad is under the reach of VAMs.

We next assessed the dynamics of dextran-rhodamine uptake in live mice, by intravital two photon microscopy (**Movie 2**). As quickly as 5 minutes after i.v. injection, eWAT macrophages were labeled with rhodamine (**Fig. 2F**).

To confirm that the uptake of dextran-rhodamine was not an aberrant result, we used a different type of molecule, the protein ovalbumin (Ova), to study the endocytic capacity of eWAT and spleen cells. Once again, VAM1 and VAM2 strongly took up fluorescently-tagged Ova, to a higher degree than splenic marginal zone macrophages (**Fig. 2G**). This association involved endocytosis and protein processing into peptides *in vivo*, as proven by the fact that whole Ovalbumin-fed VAMs were able to stimulate OTII T cells *in vitro* (**Fig. S5)**.

In summary, the gene expression profile, the location in intimate contact with the vasculature, and the high endocytic capacity of VAMs, reinforce the conclusion that, under steady state, VAMs contribute to establish and maintain a healthy adipose tissue environment.

### Most eWAT macrophages in mice fed High Fat Diet (HFD) do not have a pro-inflammatory gene expression profile

We then tested the myeloid cell populations in mice switched to a high fat diet (HFD, 60% of calories from fat, 20% carbohydrates, 20% protein), which we fed to the mice for 16 weeks (**Fig. 3A**). This treatment causes obesity and insulin resistance (Rebuffe-Scrive et al., 1993; Surwit et al., 1988). As expected, HFD induces dramatic changes in the eWAT macrophage populations (**Fig. 3B**). Confirming what has long been established (Weisberg et al., 2003; Xu et al., 2003), the total number of macrophages per gram of fat increases (**Fig. 3C**). The most apparent change is the dramatic increase in DP macrophages, which, as mentioned, express high levels of CD11b, CD11c, CD64, MHC II, and intermediate levels of CD206 (**Fig. 3B and 3C**) (Nguyen et al., 2007; Shaul et al., 2010; Wentworth et al., 2010; Zeyda et al., 2010). A very recent publication refers to these as CD9 macrophages (Hill et al., 2018), but we prefer to maintain the traditional CD11c-based DP nomenclature, as CD9 expression is proportionally more upregulated than CD11c expression in VAMs by HFD.

**Figure 3:**
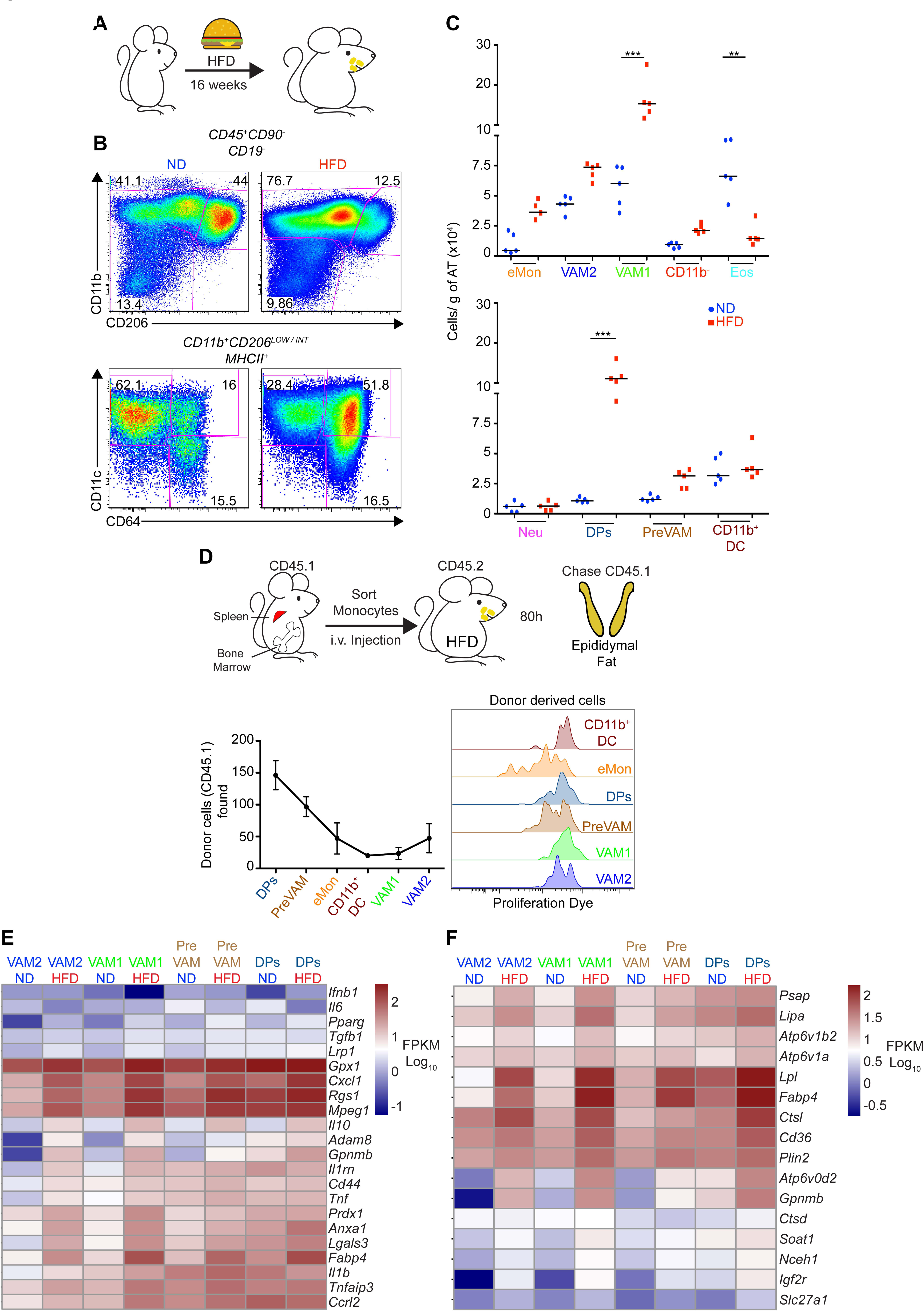
Diet-induced obesity promotes a gene signature enriched in anti-inflammatory genes in WAT macrophages. **(A-C)** WT C57BL/6J male mice 4-6 w.o. were placed on the high fat diet (HFD) OpenSource D12492i for 16 weeks, or kept on a normal diet (ND). **(B)** Flow cytometry analysis of eWAT macrophage distribution. Representative dot plot. **(C)** Myeloid cell distribution of WAT macrophages per gram of adipose tissue of ND and HFD-fed animals. Data representative of 3 independent experiments (*n*=3-5 per group). **(D)** DP macrophages are blood derived. Monocytes from bone marrow and spleen from CD45.1 WT lean mice, labeled with the Cell Proliferation Dye eFluor 450 and injected i.v. into CD45.2 HFD-fed WT mice. Eighty hours post injection, the transferred cells were quantified and characterized by flow cytometry. Representative data of 2 independent experiments (*n*=3). **(E-F)** Heatmaps of genes differentially expressed by eWAT macrophages in HFD-fed mice. **(E)** genes with anti-inflammatory and detoxification properties. **(F)** genes involved in lysosomal function and lipid metabolizing properties. **E-F** display gene expression in FPKM (Log_10_ base). *: p≤0.05. **: p≤0.01. ***: p≤0.001. Statistical analysis was carried out using unpaired T test. AT: adipose tissue.

DP are newcomer macrophages promptly generated from blood monocytes in HFD-fed mice (**Fig. 3D**). VAM1 and VAM2 also increase in number during HFD **(Fig. 3C)**, although they are very inefficiently generated by blood monocytes (**Fig. 3D**), which is consistent with the fact that VAMs are tissue-resident and are maintained, to a large extent, by self-renewal (Amano et al., 2014). PreVAMs are generated from blood monocytes, although with lower efficiency than DP macrophages, but much higher efficiency than either VAM1 or VAM2 (**Fig. 3D**). Importantly, despite the HFD-induced adipocyte hypertrophy and hypoxia (Rutkowski et al., 2015; Sun et al., 2011), none of the eWAT macrophage populations upregulated the expression of hypoxia-sensing genes, Hif1 prolyl hydrolases, or the E3 ligase that ubiquitinates *Hif1α* (**Fig. S6A**). Moreover, hypoxia regulated genes were also not upregulated by HFD feeding in eWAT macrophages (**Fig. S6A**). These results indicate that HFD-induced adipocyte stress and size expansion was not accompanied by signs of macrophage hypoxia.

Our transcriptional analysis did not confirm the view that adipose tissue macrophages arising in HFD-fed mice are overtly proinflammatory. DP macrophages arising during HFD expressed high levels of genes encoding potent secreted anti-inflammatory molecules such as IL-10 (*Il10*), TGF-β1 (*Tgfb1*) and IL-1 receptor antagonist (*Il1rn*) (**Fig. 3E**). There was no expression of α, β, or γ interferons and minimal expression of IFN-𝒦 and IL-6 in DP macrophages or VAMs before or during HFD. However, genes encoding TNFα and pro IL-1β were moderately upregulated. Many of the HFD-induced highly upregulated genes in eWAT macrophages have been previously ascribed an inflammatory signal-deactivating function, or an insulin-sensitizing function, or detoxifying functions (**Fig. 3E**). However, some of these genes, such as *Lgals3* and *Fabp4*, have been attributed with both pro- or anti-diabetogenic functions, depending on the experimental conditions, and some HFD-upregulated genes, such as *Cxcl1* and *Cd44*, are believed to exacerbate inflammation (**Fig. 3E**). A number of beneficial genes highly upregulated by HFD in eWAT macrophages are involved in lysosomal functions, as previously reported (Xu et al., 2013) (**Fig. 3F**), or in facilitating digestion of excess lipids (Aouadi et al., 2014; Singh and Cuervo, 2012)(**Fig. 3F).**

The genes highly expressed by DP macrophages from HFD-fed mice are also upregulated in VAMs (**Fig. 3E-F**). To adjust to the HFD feeding, VAMs undergo a modest increase in number but adjust their gene expression considerably. In contrast, DP macrophages experience a dramatic raise in number, but the gene expression pattern changes relatively little with HFD feeding. The few DP macrophages that exist before HFD feeding may be in the eWAT as a consequence of other injurious conditions, and thus may already have a similar pattern of gene expression. In conclusion, the gene expression pattern showed that both the newcomer CD11c^+^CD64^+^CD206^INT^ DP macrophages, as well as the pre-existing (and also expanded) CD206^HIGH^ VAMs, appear more involved in suppressing the negative consequences of HFD treatment than in the promotion or propagation of these negative effects.

### Markedly reduced endocytic capacity of eWAT macrophages from mice chronically fed HFD

Among the genes upregulated by eWAT macrophages in mice fed HFD were actin-binding proteins, such as Myosin 1E (*Myo1E*), Tropomyosin 4 (*Tpm4*), and Filamin A (*Flna*). Vimentin (*Vim*) expression was also highly upregulated (**Fig 4A**). These proteins affect the cytoskeleton of macrophages, potentially affecting cell morphology. We therefore imaged eWAT macrophages using confocal microscopy. On a normal diet, CD11b^+^CD206^HIGH^ macrophages (VAM1 and 2, which are, by far, the dominant eWAT immune cells on a ND), extend around adipocytes (**Fig 4B**, top panels). In mice fed HFD (**Fig 4B**, bottom panels), this morphology was dramatically altered. DP macrophages (CD11b^+^CD206^INT^) did not arrange themselves in the same manner. Adipocytes were bigger, and not many macrophages wrapped around them. As long ago noticed (Cinti et al., 2005), the number of crown-like structures greatly increased under HFD.

**Figure 4:**
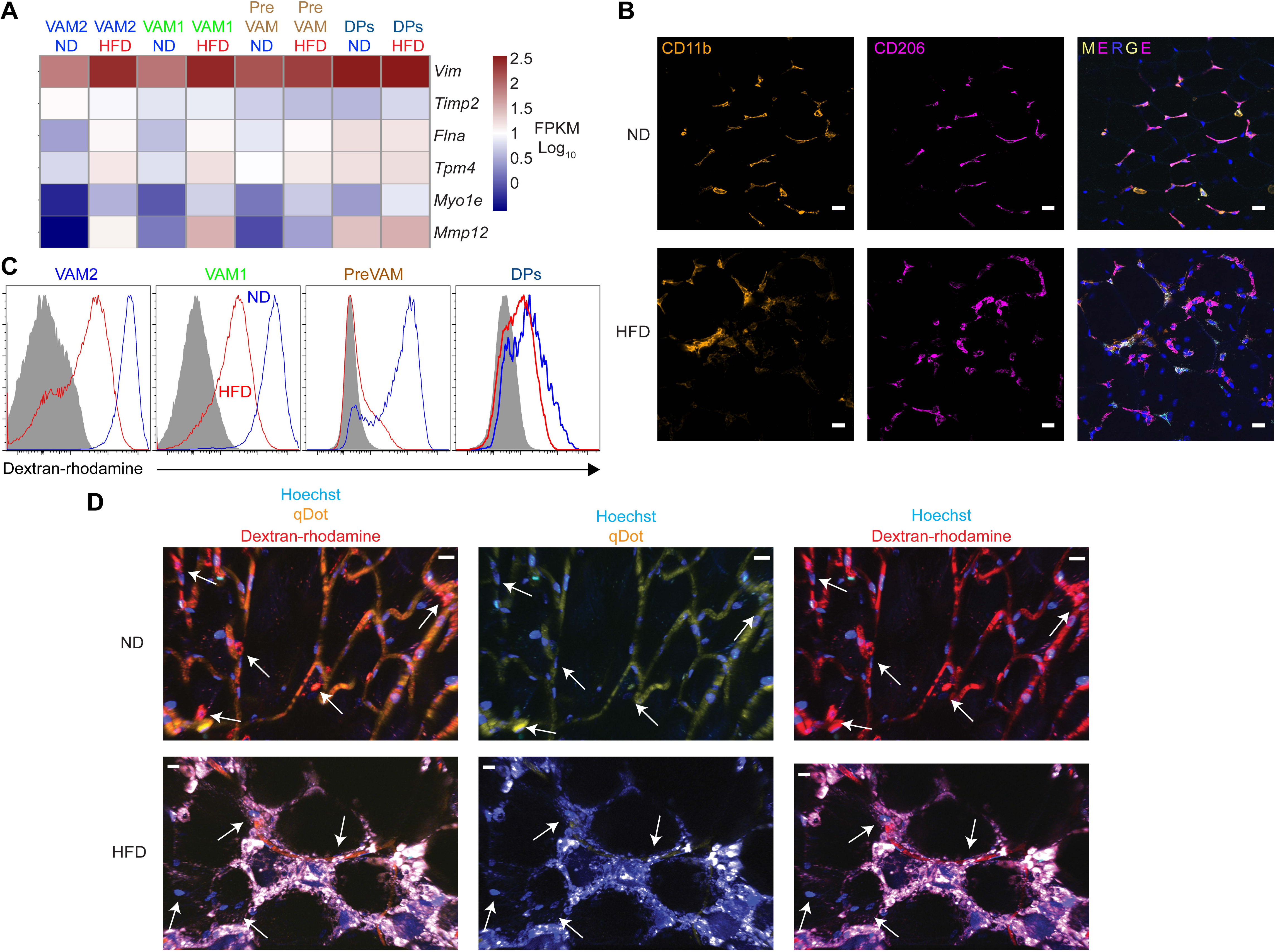
VAMs from HFD-fed mice display a diminished blood surveillance activity. **(A)** Heatmap showing genes encoding cytoskeletal proteins in eWAT myeloid cells from mice fed ND or HFD for 16 weeks. Gene expression is displayed in FPKM (Log_10_ base). **(B)** Representative confocal images of epididymal full mount sections of lean (ND) and obese (HFD) male mice showing the distribution of CD206^+^CD11b^+^ macrophages surrounding adipocytes. Scale bar =10 μm. 25x magnification. **(C)** VAM endocytic activity *in vivo* after chronic HFD feeding. dextran-rhodamine 70 kDa was injected i.v. Twenty min post-injection, animals were anesthetized and perfused intracardially. The endocytic activity of the indicated eWAT populations was quantified by the fluorescence intensity of rhodamine, as determined by flow cytometry. **(D)** Intravital microscopy analysis of eWAT dextran-rhodamine uptake in mice fed ND or HFD (**see Movie 3**). qDots (Qtracker 655), Hoechst, and dextran-rhodamine were i.v. injected and uptake of rhodamine was observed for 30 minutes. 25x magnification. Scale bar = 20 μm. Arrows in the ND-fed samples indicate VAMs uptaking rhodamine. Arrows in the HFD samples point to nucleated cells in tight proximity with blood vessels, which did not uptake rhodamine. Red labeling traces the blood vasculature.

As adipocytes become larger, surface contacts with the extracellular matrix have to be broken and re-made. DP macrophages and VAMs from HFD-fed mice robustly expressed the matrix metalloproteinase gene *Mmp12*, which could be fulfilling this function (**Fig. 4A**). Reciprocally, the expression of the gene encoding the metalloproteinase inhibitor *Timp2*, very high in VAMs under steady state, was slightly downregulated in VAMs from HFD-fed mice (**Fig. 4A**).

The reduced extent of contact between macrophages and adipocytes, and likely also between macrophages and blood vessels, made us evaluate the endocytic capacity of eWAT macrophages from HFD-fed animals. Mice were injected with dextran-rhodamine i.v., and, after 20 minutes, were sacrificed to analyze rhodamine uptake. Indeed, there was a significant reduction in the uptake of rhodamine by eWAT macrophages from HFD-fed mice, compared to ND-fed mice (**Fig. 4C**). To visualize the uptake *in vivo*, we carried out intravital microscopy, which confirmed the impaired uptake of the dye, and allowed us to visualize crown-like structures in live mice (**Movie 3** and **Fig. 4D**).

In summary, our data show that endocytic activity of eWAT macrophages in HFD-fed mice is strongly impaired, which could potentially affect their function and, thus, prolong inflammation.

### Acute inflammatory stress provokes rapid and reversible changes in eWAT macrophage populations

Having addressed the response of eWAT myeloid cells during chronic inflammatory conditions, we next studied the impact of acute inflammation on the composition of the different myeloid eWAT cell populations (**Fig. 5A**). 24 h after i.v. administration of 50μg LPS, a sublethal dose, the macrophage populations were dramatically changed in frequency and absolute numbers. VAMs 1 and 2 were rapidly reduced by approximately 4-fold, and the number of DP macrophages (CD64^+^ CD11c^+^) more than tripled (**Fig. 5B and 5C**). These changes were already significant at 12 h (**Fig. 5C**). The number of CD11b^-^ dendritic cells did not change upon acute LPS-driven inflammation, but the number of CD11b^+^ dendritic cells was greatly reduced within 12 h. Monocytes and neutrophils were increased. Almost all myeloid cell populations returned to the steady state composition by day 5 (**Fig. 5C**). Although published data and our experiments shown in **Fig. 3D** indicate that VAMs and DP macrophages probably have a different origin, the fact that VAMs decreased rapidly, and, at the same time, DP macrophages increased, made us investigate the possibility that LPS could induce the conversion of VAMs into DP macrophages. To investigate this possible conversion, we used a “functional fate mapping” strategy. We took advantage of the very high endocytic capacity of VAMs, to label them with dextran-rhodamine and follow the fate of labeled cells. We injected dextran-rhodamine i.v. and, 36 hours later, injected LPS i.v. 24 h after LPS injection, we analyzed the eWAT myeloid cells by FACS (**Fig. 5D**). As expected, VAMs efficiently took up dextran-rhodamine, which remained in these cells after LPS treatment (**Fig. 5E**). Importantly, the DP macrophages were not labeled with rhodamine, indicating that they did not derive from converted VAMs (**Fig. 5E**). Most likely, they are derived from circulating monocytes, similarly to the equivalent DP population that robustly expands in the eWAT during HFD feeding. In contrast to VAMs, preVAMs lost their rhodamine after LPS treatment, suggesting that they were quickly replaced by unlabeled cells.

**Figure 5:**
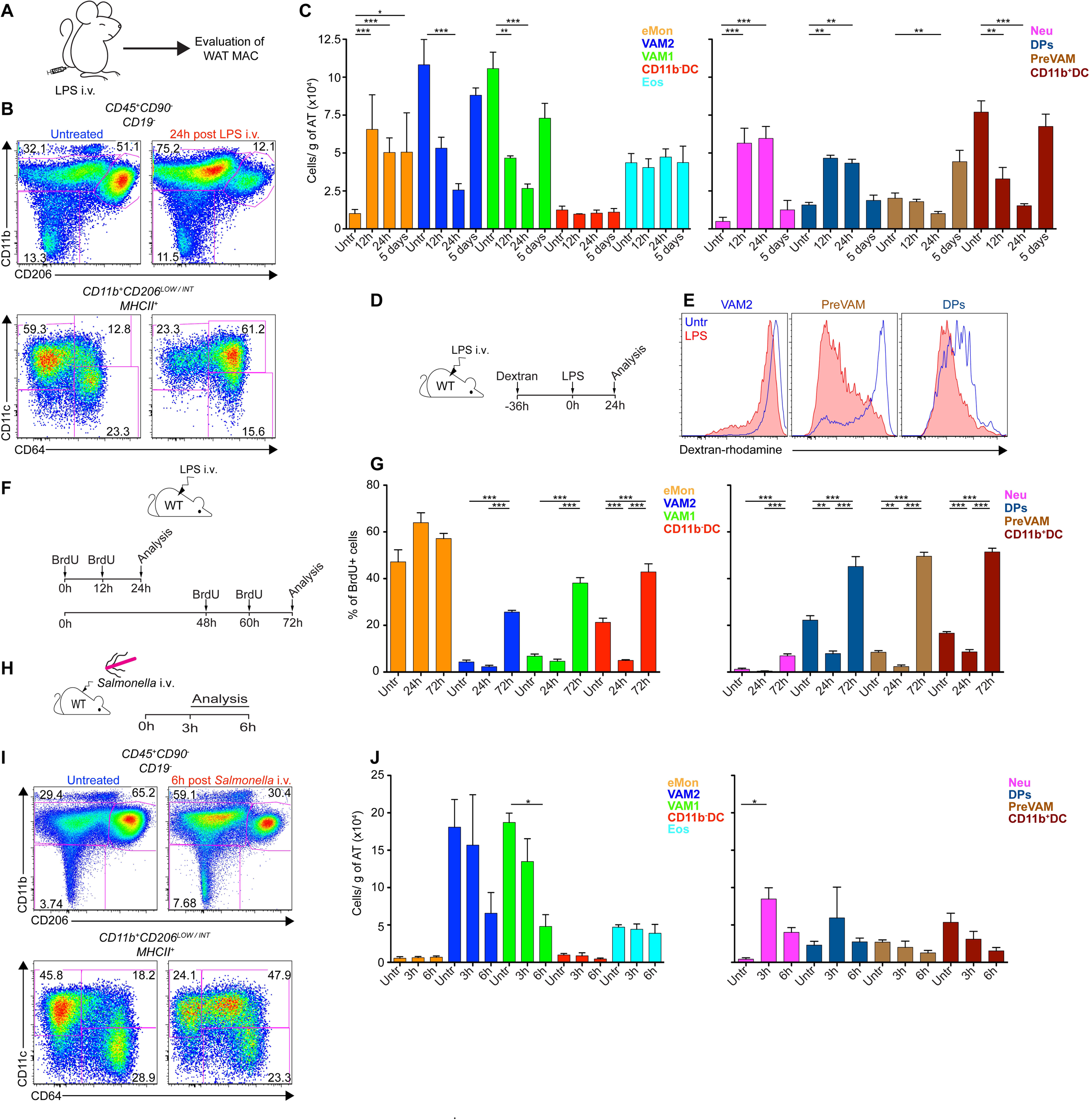
Differential response of the eWAT macrophage populations to systemic acute inflammatory challenges. **(A-C)** WT mice were injected 50μg of LPS i.v., and the eWAT myeloid cell composition was determined 12, 24 and 120h post injection. **(B-C)** Flow cytometry **(B)** and absolute number **(C)** analysis of the eWAT myeloid compartment. Representative data of 3 independent experiments (*n*= 3-5 mice). **(D-E)** Functional fate mapping of VAMs exposed *in vivo* to LPS. WT mice were injected i.v. with dextran-rhodamine 70 kDa. Thirty six hours later, the same animals were injected with 50μg of LPS. 24h after the LPS injection, eWAT macrophages were analyzed by flow cytometry. Data representative of 2 independent experiments (*n*=3 per group). **(F-G)** eWAT myeloid cells proliferation determined by BrdU incorporation at different time-points. WT animals were injected i.p. twice with 1mg of BrdU 12h apart. The BrdU injections started concomitantly with LPS injections, or 48h post LPS injection. Animals were analyzed at 24h or 72h time points, respectively. Representative data from 2 independent experiments (*n*=3-5 per group). **(H-J)** WT mice were inject i.v. with 10^8^ CFU of *Salmonella enterica* serovar Tymphimurium (*Salmonella*) and eWAT myeloid cell composition was determined 3 and 6h post infection. **(I-J)** Flow cytometry and absolute number of eWAT myeloid cell populations. Representative data of 2 independent experiments (*n*= 3-5 mice). *: p≤0.05. **: p≤0.01. ***: p≤0.001. Statistical analysis was carried out using one way anova together with turkey post test. AT: adipose tissue.

If the rapid reduction in VAM numbers was due to a rare emigration of mature tissue macrophages from eWAT, one of the possible alternative locations would be the spleen. We therefore also determined the myeloid changes in the spleen 24h after i.v. LPS injection. Macrophages with the surface profile of eWAT VAMs (CD11B^+^CD206^HIGH^CD64^HIGH^) were not found, indicating that they did not migrate from the eWAT to the spleen. The number of splenic marginal zone macrophages was not significantly changed. As expected, neutrophils were highly increased. Monocytes were sharply decreased in the spleen, and eosinophils and CD11b^+^DC were decreased as well (**Fig. S7A and S7B**). Monocytes were increased by LPS in the eWAT, and eWAT DP macrophages, which are most likely derived from monocytes, were increased in the eWAT as well, perhaps providing a balance between monocytes in tissues and those in lymphoid organs.

To evaluate the contribution of apoptotic cell death on the eWAT subpopulation reshaping after LPS treatment, we determined the expression levels of active caspase 3, the main executioner caspase, by flow cytometry, but we did not observe an increase in any eWAT myeloid population (**Fig. S7C**). We noted that, at baseline, both DC populations, CD11b^+^ and CD11b^-^ DC, display high levels of activated caspase 3. Thus, LPS causes rapid changes in the eWAT myeloid cells, with a reduction in VAMs that we cannot ascribe to differentiation into a different cell type, migration to the spleen, or caspase-3-mediated apoptosis.

As the number (and proportion) of eWAT VAMs is rapidly and dramatically reduced by LPS, it is equally striking that the eWAT VAM populations returns to steady state numbers by 5 days post LPS administration. Given the fact that WAT macrophages have been shown to divide locally (Amano et al., 2014), we expected that, after LPS, the repopulation of VAMs would take place by their own cell division. We therefore monitored cell division by BrdU incorporation over the first and third days of the experiment (**Fig. 5F**). Without LPS, more than half of eWAT monocytes, expectedly, become BrdU^+^ during the 24 h labeling (**Fig. 5G**). At a 20% BrdU plateau are the two DC populations, indicating a relatively fast turnover. VAMs 1 and 2 incorporate a very low amount of BrdU. Among macrophages, DP and preVAMs, the two populations that we showed to be monocyte-derived, are the only populations with a measurable BrdU incorporation (**Fig. 5G**). Upon LPS treatment, BrdU incorporation into VAMs remained very low during the first day, but increased at the third day (**Fig. 5G**). This is the time in which the VAMs begin their recovery from LPS. This result is compatible with the self-division of VAMs that was discussed above, but it does not exclude that dividing monocytes took up BrdU and very rapidly differentiated into VAMs.

To assess whether bacterial infections could also cause a rapid alteration in the composition of the eWAT, we infected mice with the Gram-negative bacteria, *Salmonella enterica* serovar Tymphimurium (hereafter *Salmonella*) (**Fig. 5H**). *Salmonella* septicemia is a dangerous complication of *Salmonella* infections, and to study this sequela we injected the bacteria i.v. Similarly to LPS, systemic *Salmonella* infection rapidly caused a reduction of VAMs by 3 and 6 h (**Fig. 5I and 5J**). Also similarly to LPS, *Salmonella* caused an increase in DP macrophages, although it was of shorter duration than the increase caused by LPS, and lower in magnitude (**Fig. 5J**). Neutrophils were also increased, but, in contrast to LPS, we did not observe an increase in eWAT monocytes, at least at the early time points that we studied (**Fig. 5J**). However, in the spleen, *Salmonella* causes a rapid monocyte reduction, which is similar to that observed with LPS (**Fig. S7D and S7E**).

Thus, the common feature of LPS and Gram negative bacterial infection is that both induce a very rapid reduction in number and frequency of eWAT VAMs, with increase in DP macrophages. Alterations in eWAT myeloid cells were not, however, limited to LPS and Gram-negative bacteria. Infection with the Gram-positive bacteria *Staphylococcus aureus* (hereafter S. aureus) also resulted in a decrease in the number of VAMs (**Fig. S7F-H**), more markedly in VAM1 (**Fig. S7H**). Analogous to LPS administration, S. aureus infection caused an increase in the proportion of eWAT monocytes (**Fig. S7H**). In the spleen, there was a major difference between S. aureus infection and LPS or *Salmonella* infection: S. aureus caused a marked increase in monocytes at 12h post-infection (**Fig. S7I and J**).

We conclude from these data that LPS or systemic bacterial infections have an immediate impact on the composition of eWAT myeloid cells, in particular causing a reversible decrease in VAMs.

### Fasting reduces the number of eWAT Vasculature-associated Adipose Macrophages

Given the rapid fluctuation of VAMs and DP macrophages upon acute inflammatory signals, we studied a model of acute nutritional stress, provoked by 24h fasting (**Fig. 6A**).

**Figure 6:**
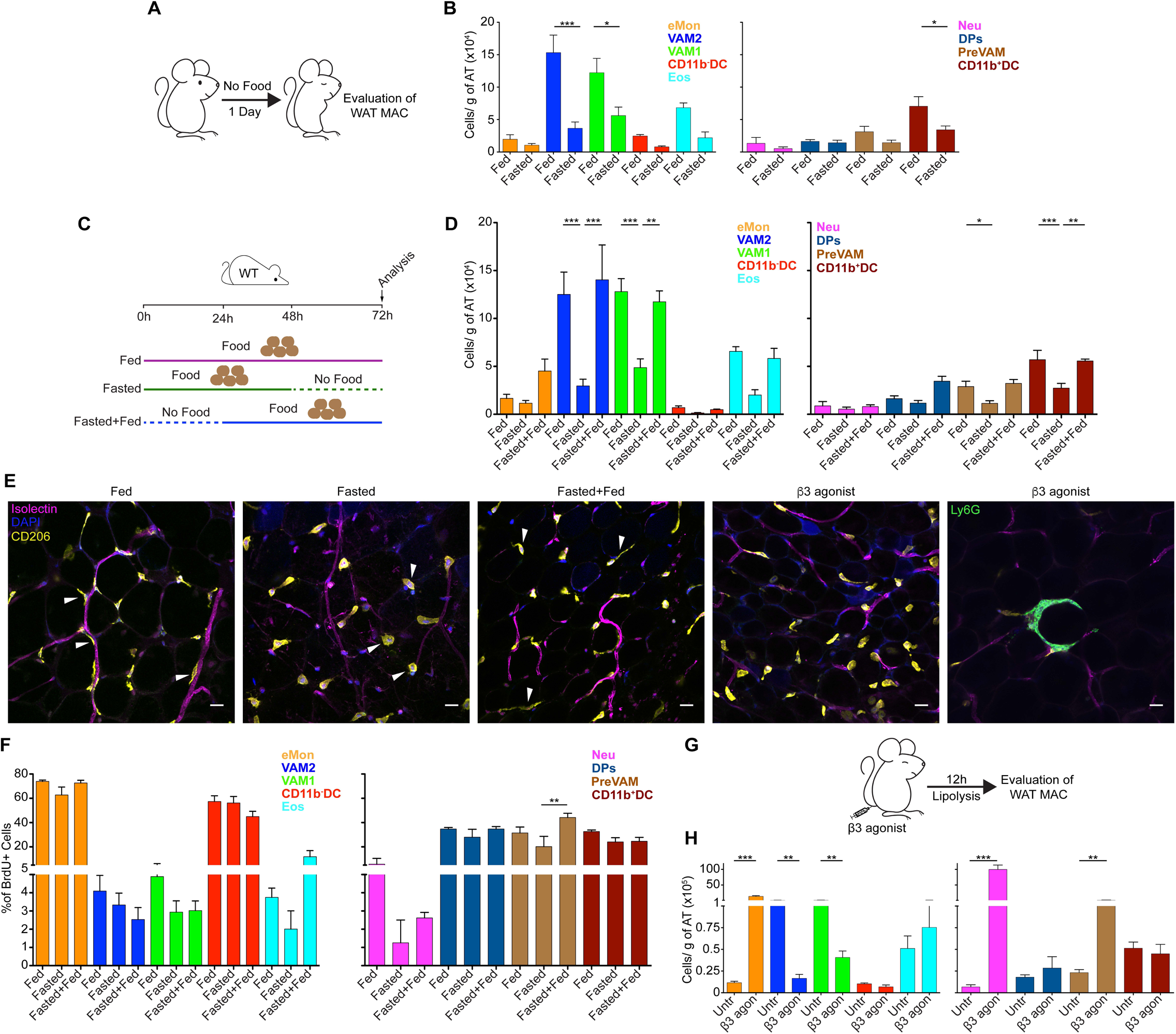
Fasting swiftly reduces the number of eWAT VAMs. (A-B) WT mice were fasted for 1 day and the distribution of eWAT myeloid cells evaluated by flow cytometry. **(A)** Experimental scheme. **(B)** Absolute number of eWAT myeloid cells. Representative data of 3 independent experiments (*n*= 3-5 mice per group). **(C-D)** Recovery of eWAT VAM numbers after food was restored. WT mice were subjected to fasting for 1 day (Fasted), to fasting for 1 day and recovery for 2 days (Fasted+Fed) or were never deprived of food (Fed), as schematically shown in **C**. **(D)** Myeloid cell population numbers in the eWAT. Representative data of 2 independent experiments (*n*= 3-5 mice per group). **(E)** Representative confocal images of epididymal full mount sections of WT mice fed, fasted, fasted+fed or treated with the β3 adrenergic receptor agonist CL 316243. Pictures shows the distribution and morphology of VAMs (Arrowheads. CD206^+^DAPI^+^) around adipocytes and blood vessels, labeled with Isolectin. The far right panel shows an atypical crown-like structure formed by accumulation of neutrophils (Ly6G^+^) around an adipocyte. Scale bar =10 μm. 25x magnification. **(F)** Proliferation of the different myeloid cell populations shown in **Fig. 6C-D** as determined by BrdU incorporation. 1mg of BrdU was injected twice i.p. 12h apart in the last 24h of the experiment. Representative data from 2 independent experiments (*n*=3-5 per group). **(G-H)** Lipolysis was induced in WT mice by the i.v. injection of β3 adrenergic receptor agonist CL 316243, without food withdrawal. Twenty hours post-injection, the distribution of myeloid cells was assessed as indicated above. MAC: macrophages. Representative data from 4 independent experiments (*n*=3-5 per group). *: p≤0.05. **: p≤0.01. ***: p≤0.001. Statistical analysis was carried out using one way anova together with turkey post test or unpaired T test (only for pair comparisons).

Upon food withdrawal, blood glucose levels decrease and lipolysis is triggered, which results in the release of non-esterified fatty acids (FFA) into the serum. As fasting continues, FFA uptake from the blood by cells such as skeletal muscle cells also increases. To test parameters in mice under our conditions, we measured FFA, blood glucose, and weight loss during a 24h fasting experiment. Serum FFA concentration peaked at approximately 12-16h of fasting, indicating a net lipolysis and release of FFA, and thereafter serum FFA concentration diminished very slightly (**Fig. S8A and B**). As expected, blood glucose decreased early (**Fig S8C**). After 24h, body weight loss was approximately 10% (**Fig. S8D**). Strikingly, myeloid cells were dramatically reduced by 1d fasting. VAM numbers were the most affected by fasting. However, not only VAMs were reduced; also preVAMs, eosinophils, and both CD11b^+^ and CD11b^-^ DC were reduced (**Fig. 6B**). Unlike HFD and LPS, fasting did not provoke an increase in the DP macrophage population; instead, it dropped, albeit non-significantly. Similarly to what we observed after LPS administration, we did not detect a surge in active Caspase 3 concomitantly with the rapid cell decrease (**Fig. S8E**).

In nature, mice must endure fluctuations in the food supply. Given the striking reduction of VAMs that we observed after one day of fasting, we surmised that the number of VAMs would rebound in rapid fashion; otherwise, in nature mice would always have low eWAT VAMs. We therefore placed a group of one-day fasted animals back on normal diet, and evaluated the myeloid eWAT populations after two days back on normal diet (labeled: fasted+fed) (**Fig. 6C**). To have the appropriate controls in the same experiment, we had a group of mice that was normally fed, never fasted (labeled: fed), and a group of mice that was fasted the last day (labeled: fasted) (**Fig. 6C**). As expected, animals feeding after fasting quickly re-established their initial weight (**Fig. S8D**). Remarkably, after 48h of re-feeding, the eWAT populations were back to the normal situation (**Fig. 6D**). To evaluate the VAM morphology after fasting, we also carried out a confocal microscopy analysis. Confocal images of the eWAT showed that, compared to mice fed all along, VAMs 1 and 2 (CD206^HIGH^) from 1d fasted mice were abnormally rounded in shape (arrowheads), indicating that they partially lost their contacts with vasculature and adipocytes (**Fig. 6E**). While we chose a microscope field that contained many VAMs to allow the morphological characterization, the reduction in VAM numbers was also noticeable by our confocal analysis. The contorted vasculature, which surrounds adipocytes, showed the expected reduction in adipocyte diameter upon fasting. eWAT VAMs from fasted+fed mice, were abundant in number, but had only incompletely re-gained their normal topology (**Fig. 6E**). Thus, the VAM depletion induced by 1d fasting is brief and, it appears, fully reversible.

To assess whether the reconstitution of eWAT VAM numbers was due to self-division, we monitored the incorporation of BrdU over the last 24h of the fasting and re-feeding experiment described above. This gives us a fed (baseline) group, a 24h fasted group, and a group that was fasted and had fed over the last 2d. BrdU was not incorporated by VAMs either during the day of fasting, or during the second day of re-feeding, when the cells would be expected to be actively dividing (**Fig. 6F**). Most likely, VAMs were generated by monocytes or preVAMs that had divided before the BrdU pulse, and were differentiating to VAMs while the BrdU pulse was ongoing. It thus appears that, to rapidly reconstitute VAMs after fasting, mice made use of monocytes, or monocyte-derived preVAMs.

Fasting causes a series of metabolic changes, of which increased lipolysis is one of them. It was possible that the eWAT VAM reduction was directly caused by lipolysis, with remodeling of the niche that VAMs occupy in steady state, or in response to other metabolic parameters, such as lowering of glucose, insulin or the like. To induce lipolysis without food withdrawal, we administered a β3 adrenergic agonist, CL 316,243 (Yoshida et al., 1994). Adipocytes express β3 adrenergic receptors, which are locally triggered by sympathetic nerve activation, or systemically by adrenal catecholamines (Bartness et al., 2014). Like fasting, CL 316,243 also induced a dramatic drop in VAM populations after 12h of a single 2mg/kg dose (**Fig. 6G-H**). Intriguingly, there was a major influx of monocytes and neutrophils into the eWAT (**Fig. 6H, Fig. S8F-H**). By confocal microscopy, we could clearly observe that CL 316,243 reduced and changed the morphology of CD206^HIGH^ (VAMs) macrophages, but induced crown-like structures that are atypical, as they are composed of Ly6G^+^ neutrophils, not macrophages (**Fig. 6E**). In sum, both fasting and a β3 adrenergic agonist caused the rapid depletion of VAMs in the eWAT, suggesting that lipolysis itself, and not the nutritional state, is the driving force in the VAM reduction.

In conclusion, our results show adaptation of adipose tissue microphages to diverse environmental challenges.

## Discussion

In this report, we characterized the behavior of the different eWAT macrophage populations in response to chronic and acute inflammatory and metabolic stress. By gene expression profiling, flow cytometry and imaging, we could distinguish four populations of macrophages, which we referred to as Vasculature-associated Adipose Macrophages 1 and 2, or, in short, VAM1 and VAM2, PreVAM, and CD64^+^CD11c^+^ Double Positive (DP) macrophages. In steady state, by far VAM1 and VAM2 are the two dominant populations. VAM1 and VAM2 are similar in gene expression, and very likely to exert analogous functions. Previous characterization of adipose tissue macrophages (ATM) in healthy mice undoubtedly focused on these dominant macrophages. Although VAM1 and VAM2 poorly express two of the more typical M2 macrophage genes, Arginase 1 (*Arg1*) and Chitinase-like 3 (*Chil3*), the overall expression pattern resembles M2 macrophages, as they express high levels of CD206, CD301a, CD163, CD209, and *Retnla*/Fizz1. They express genes that allow the uptake of apoptotic cells, many scavenger receptors, antioxidant and detoxifying molecules. VAMs from lean mice harbor abundant lipid droplets, which had been reported in adipose tissue of obese mice (Aouadi et al., 2014). The most previously underappreciated aspect of the VAMs is their very close contact with vessels, leading to a high endocytic capacity of blood molecules. Their high endocytosis, gene expression and abundant lipid droplets indicate that, in mice fed a normal diet, VAM1 and VAM2 are not passively waiting for disturbance, but rather, they appear very active under steady state. VAM properties suggest that a possible function of these tissue-resident macrophages is to keep clean the eWAT adipose tissue environment, both by removing noxious catabolites from the adipocytes, and by forming a barrier from blood toxins. A correlation between phagocytosis capacity, CD206 expression and tissue homeostasis by macrophages in different organs has recently been reported (N et al., 2017).

Our work shows no evidence of an M2 to a Classically Activated/M1 shift during diet-induced obesity (DIO). The initial M1/M2 paradigm was drawn on the competing activities of Arginase (Arg) and Nitric Oxide Synthase (Nos) for Arginine. When Arg is highly expressed, no or little NO can be generated, changing the biological properties of the cells and their environment (Mills et al., 2000; Rath et al., 2014; Rodriguez et al., 2017). However, before and after the onset of diet-induced obesity eWAT macrophages lack expression of both Arg and iNOS (*Nos2*). Most clearly, DIO does not induce a big switch to M1-like macrophages. In fact, among the genes prominently expressed by DP macrophages after HFD feeding was CD180, which was described in a screen for IL-4-induced transcripts in macrophages (Czimmerer et al., 2012). *Egr2*, another M2 gene, was also upregulated, while *Cd38*, defined as an exclusive M1 marker (Jablonski et al., 2015), was strongly downregulated in VAMs by HFD feeding (**Fig. S6B**). Furthermore, we monitored the expression of genes that characterize murine macrophages stimulated with LPS or IFNγ (Murray et al., 2014). None of these characteristic M1-like genes, such as *Ido1, Ido2, Marco, Il27*, IL-23 p19, IL-12p40, IL-12p35 and *Socs1* are significantly expressed by eWAT macrophages, under normal or high fat diet (**Fig. S6B)**. *Nfkbiz* and *Irf5* are, indeed, expressed by the HFD-induced monocyte-derived DP macrophages, but they are also expressed at very similar levels by VAMs under normal diet **(Fig. S6B)**, and even by eWAT monocytes and dendritic cells. A metabolic shift was noted in M1 macrophages, whereby pyruvate gets carboxylated to oxaloacetate by the enzyme pyruvate carboxylase (*Pcx*), a possible pathway to gluconeogenesis or to restore oxaloacetate levels. However, *Pcx* is not expressed by any macrophage type under HFD. It has been proposed that Group 1 ILC (Gr-1 ILC) produced IFNγ to drive M1 ATM differentiation in the adipose tissue (O’Sullivan et al., 2016). Gr-1 ILC expand during HFD treatment, presumably driving M1 conversion. Blocking IFNγ or eliminating Gr-1 ILC with anti-NK1.1 increased the M2/M1 ratio and restored insulin sensitivity (O’Sullivan et al., 2016). However, these M1 macrophages were characterized by their expression of CD11c, not by the expression of a set of genes traditionally associated with M1 macrophages. Our data also show that HFD triggers a massive inflow of monocyte-derived DP macrophages, which express CD11c, although they are not M1 or M1-like. It was also reported that Gr-1 ILC could kill ATMs during steady state, with preference for M2 macrophages, but this cytotoxic capacity is impaired in mice fed HFD (Boulenouar et al., 2017). In that report, M1 macrophages were also defined by their expression of CD11c, which, as indicated above, we believe is inaccurate.

One of the possible explanations for the M1 discrepancy is that, most frequently, qPCR was used to compare the macrophage populations pre- and post-HFD exposure. However, robust fold induction by qPCR could be the result of a very small minority of M1 cells. In that regard, RNAseq data can better estimate the transcript levels of a cell population.

Our data are more compatible with the category of “Metabolically-activated” macrophages (MMe), discovered by stimulating M0 macrophages *in vitro* with a mixture of insulin, glucose and palmitate (Kratz et al., 2014), and subsequently validated *ex vivo*. The MMe activation is independent of pro-inflammatory signals (Kratz et al., 2014). Among the key MMe genes, *Cd36* and *Plin2* are highly upregulated in our eWAT samples by HFD, and thus in good agreement with the MMe nomenclature. However, *Abca1* expression is not upregulated, although it is already very high in VAMs from lean mice, and, in addition, we did not observe upregulation of *Acsl1, Bag6, Dhcr7, Far1, Plxnb2* or *Ptprj* (**Fig S6B**).

The vast majority of genes highly expressed by DP macrophages, which in terms of fold increase is the population most boosted by HFD feeding, also became highly expressed by VAMs when mice are switched to HFD. For example, VAMs increased their expression of the IL-10, TGF-β1, and IL-1 receptor antagonist (*Il1rn*) genes. In fact, we know of no gene whose expression is highly upregulated in DP macrophages and decreased in VAMs in mice fed HFD, or vice-versa. VAMs displayed a reduced expression of the M2 genes described above upon HFD feeding; this includes *Mrc1*, the gene encoding CD206, which is slightly downregulated by HFD. These adaptations reflect the high environmental pressure placed on the visceral adipose tissue by chronic HFD treatment.

Over the past decade, it has become apparent that many tissue-resident macrophages are not derived from bone marrow progenitors, via circulating monocytes, as it had previously been thought (Ginhoux and Guilliams, 2016; Schulz et al., 2012). Typical examples of self-renewing non bone marrow-derived macrophages in adult animals are microglia and Langerhans cells (Ajami et al., 2007; Chorro et al., 2009; Ginhoux et al., 2010; Merad et al., 2002). In the adipose tissue, self-renewal of macrophages was also shown to be the case (Amano et al., 2014). Our, data tested this concept in a number of pathophysiological conditions. VAMs in the eWAT of healthy mice have, indeed, characteristics of tissue-resident macrophages. However, our work showed that they adjust very rapidly to changes in nutrition and inflammation status. Under acutely stressful conditions, the number and proportion of VAMs decreases substantially, and returns to normal shortly after the stressful condition is removed. In the case of fasting, the reconstitution of VAM numbers appears to be relying on dividing monocytes.

PreVAMs have a gene expression signature that is quite similar to that of VAMs (**Fig. 1B**). They also display intermediate endocytic capacity to that of VAMs. We showed that preVAMs are generated from monocytes in mice fed HFD, yet their number does not increase nearly as much as the number of DP macrophages, which are also efficiently generated by monocytes. After LPS challenge, at the same time as VAMs were depleted, preVAMs lost much of their incorporated rhodamine, while VAMs did not (**Fig. 5E**). This suggests that, under these conditions, preVAMs are rapidly renewed. We propose that, under acute nutritional stress, preVAMs or monocytes can replenish VAMs, which, under physiological or mild inflammatory conditions, are able to maintain their number by cell division, without major input from other populations.

It is believed that macrophages play a pathologic role to mice during HFD treatment. CD11c-DTR (diphtheria toxin receptor) have been used to ablate all CD11c^+^ cells after diphtheria toxin (DT) administration, including but clearly not limited to fat DP macrophages. This treatment resulted in a reduction of HFD-induced adipose macrophages and crown-like structures, and, importantly, caused an improvement of glucose and insulin sensitivity (Patsouris et al., 2008). However, the authors’ data also show that DT administration had a beneficial impact on mice that did not express the DTR. Given our observations that VAMs are highly associated with the vasculature and highly endocytic, it is possible that diphtheria toxin could be taken up by macrophages that did not express CD11c, thus confounding the interpretation of the authors’ data. One major additional caveat of the CD11c DTR experiment is that the removal of DC would deactivate effector T cells, which could be responsible for the observed improvement in insulin sensitivity.

There is a rapid reduction of eWAT VAMs upon acute stress caused by i.v. LPS or bacterial infections with *Salmonella* or S. aureus. We determined that the VAMs were not converting to other eWAT cell types, that they did not migrate to the spleen, and we did not observe signs of Caspase 3-mediated apoptosis. Several Caspase 3-independent programmed cell death pathways have been reported in the literature (Lee et al., 2006; O’Sullivan et al., 2007). It has been long known that, *in vitro*, LPS or LPS-containing bacteria can cause apoptosis of macrophages (Bingisser et al., 1996; Hilbi et al., 1997; Xaus et al., 2000), but it is also known that LPS is a powerful activator of macrophages, through TLR4 signaling. It has been shown that Gr-1 ILC could kill adipose tissue macrophages (Boulenouar et al., 2017), and that they do it more efficiently with M2 macrophages, such as VAMs. All myeloid cells in the eWAT express equivalently low levels of *Rae1* mRNA, a target of NKG2D, regardless of the diet. However, we did not determine whether rapid increases in *Rae1* expression levels occurred after LPS, *Salmonella* or S. aureus infection, or fasting. It is therefore possible that Gr-1 ILC are the executioners of VAMs very rapidly after acute infectious or metabolic stress.

We studied the spleen as a place where eWAT VAMs could migrate, because of its strategic location in the circulation. Given that we cannot account for either apoptotic or further differentiated VAMs within the eWAT, we cannot rule out that live eWAT macrophages could enter the vasculature and end up in places other than the spleen, such as draining nodes, liver, lung or bone marrow. However, it would be unlikely for mature tissue-resident macrophages such as VAMs to circulate out of the home organ.

Bacterial infections (or LPS) cause metabolic changes in the host, which in turn may be associated with rapid changes in adipose myeloid cell composition. Infections induce “sickness behavior”, whereby the host reduces food consumption, triggering gluconeogenesis and lipolysis, and a higher brain consumption of ketone bodies (O’Neill and Hardie, 2013; Wang et al., 2016). LPS itself was shown to increase glucose uptake and aerobic glycolysis in macrophages (Krawczyk et al., 2010), mediated by the pyruvate kinase M2 isoform (Palsson-McDermott et al., 2015). Thus, the rapid and reversible reduction of eWAT VAMs caused by *Salmonella* or LPS could represent a response similar to the fasting-induced reduction of eWAT VAMs that we uncovered in this manuscript.

In contrast to the dominance of VAM1 and 2, DP macrophages represent only 1/6 of the VAM1 and 2 macrophages under steady state conditions. This DP macrophage population is dramatically increased by both chronic HFD and acute LPS i.v. injection (**Fig. 3B-C and 5B-C**). However, beyond sharing the key surface markers used to define DP macrophages (**Fig. 1A and S1L**), we do not know the degree of similarity between these LPS- or *Salmonella*-induced DP macrophages and the HFD-induced DP macrophages.

In the life of wild mice, fluctuations in dietary intake are to be expected. In this context, the reduction in eWAT VAMs, eosinophils, monocytes and DC after just one day without food is remarkable. Steady state numbers are, however, rapidly restored upon normalization of the food intake. Fasting-induced lipolysis provides circulating non-esterified fatty acids, which can be used by other cells (e.g. skeletal muscle cells, hepatocytes) to provide energy. During this fasting period, the liver is known to activate gluconeogenesis in response to ghrelin, thus maintaining blood glycaemia at acceptable levels. Lipolysis causes, naturally, adipocytes to shrink (**Fig. 6E**), and we speculate that this change in size removes cytoskeletal cues from the VAMs, possibly causing a number of them to undergo cell death. Thus, adipose tissue remodeling induced the loss of VAMs. The effect of fasting on eWAT macrophages was mimicked by a β3 adrenergic agonist, CL 316243, in conditions of food availability. These results support the view that lipolysis itself, not the nutritional state, caused the loss of VAMs.

Interestingly, another non-infectious manipulation of adipose tissue, which is the induction of adipocyte death by overexpression of an active Caspase 8, was reported to cause an increase of macrophages that displayed an M2 activation profile, rather than the reduction that we observed with fasting (Fischer-Posovszky et al., 2011). Caspase 8, however, induces death of adipocytes, and these apoptotic cells require clearance by macrophages. Lipolysis, on the other hand, changes the architecture of the adipose tissue by reducing the size of the cells to which VAMs are attached, without massive apoptosis.

Lipolysis is not only induced under energy needs, but it is known that insulin-resistant obese animals have an increased lipolytic activity and serum concentration of FFA (Duncan et al., 2007), presumably to help mitigate the cellular stress associated with adipocyte hypertrophy. However, this is known to be triggered by different stimuli than fasting-induced lipolysis (Duncan et al., 2007; Zechner et al., 2009).

The strong inflammatory response to a single CL 316,243 dose was not unexpected. Acute inflammation occurred using both 2mg/kg and 0.2mg/kg doses. CL 316,243 is a small compound, which is endotoxin-free. It was reported that one i.p. treatment with CL 316,243 resulted in an increased myeloid population in the WAT after 6h (Mottillo et al., 2007). Chronic CL 316,243 treatment with osmotic pumps was reported to cause rapid inflammation in the adipose tissue, which was more marked in Pparg-deficient than WT mice (Li et al., 2005). Kosteli and coworkers fasted mice for 24h and quantified the mRNA expression of *Adgre1* (F4/80) and *Csfr1* (CD115) by qPCR, and concluded that eWAT macrophages had increased; they also administered CL 316,243 i.p. and observed an increase in F4/80 staining by immunohistology after 18h (Kosteli et al., 2010). We have carefully characterized the myeloid infiltrating cells by flow cytometry and confocal microscopy after fasting and i.v. CL 316,243 treatment, and established that VAMs were significantly decreased in both situations, DP macrophages were not significantly changed, and monocytes and neutrophils were increased, but only by CL 316243. A possible explanation for the lack of neutrophilia after fasting, is that besides lipolysis, CL 316243 induces thermogenesis (Kazak et al., 2017), which fasting for one night does not, and this could provide the basis for the transient inflammatory response that we observed. β3 adrenergic agonists are currently FDA approved for human use in overactive bladder syndrome (Malik et al., 2012; Takasu et al., 2007), and have been discussed as anti-obesity and insulin-sensitizing treatments for humans (Arch, 2008; Cypess et al., 2015). It is therefore important that the basis of the pro-inflammatory response caused by β3 adrenergic agonists be further studied.

A major unresolved issue raised by our work is the possible adaptive value of acutely removing potentially valuable macrophages such as VAMs during inflammatory or nutritional stress. During acute infections, it is likely that the monocyte-derived DP macrophages are better equipped than VAMs to participate in pathogen removal functions. It is less clear what the adaptive value of transiently losing VAMs is during fasting, if there is any. Experiments to address this question directly, such as injecting VAMs to repopulate the WAT are technically very difficult. The natural depletion of VAMs is also likely to affect the integration with other immune cells such as effector and regulatory T cells, which abound in white adipose tissue. Thus, our characterization of the several different myeloid cell types could help the clarification of the interactions between lymphoid and myeloid cells in adipose tissue.

In conclusion, we showed how adipose tissue macrophages, which are located at the intersection between the vasculature and the adipose tissue, are very sensitive to different environmental changes, yet maintain a striking capacity for rapid recovery.

## Materials and Methods

### Mice

Foxp3^GFP^ male mice in C57BL/10.PL background (Weiss et al., 2012) and C57BL/6J (000664) purchased from Jackson laboratories were used as WT animals along the study. B6 CD45.1 (564) animals were purchased from Charles River. CX3CR1-GFP mice (005582) were purchased from Jackson laboratories. High-fat diet (HFD)-fed male mice were treated with a HFD for at least 16 weeks starting at 4-6 weeks of age. Mice were fed standard chow providing 17% calories from fat (LabDiet Formulab 5008) or a HFD providing 60% calories from fat (Research Diets, D12492i). Blood glucose was determined with a One Touch Basic glucometer (Lifescan, Milipitas, California). Mice were sacrificed by CO2 euthanasia and serum was collected when necessary. Serum analysis of non-esterified fatty acids (FFA) content was performed using a FFA assay kit (Abcam, ab65341). As indicated in the text, some mice were treated i.v. with 2 mg/kg of β3 receptor agonist (CL 316243, Tocris-1499), 50 μg of LPS (Sigma-Aldrich, L4391), 18mg/kg of dextran-rhodamine 70 kDa (Sigma-Aldrich, R9379), 2mg/kg of OVA-A647 (Thermofisher, O34784). Animals were housed at NYU Medical Center Animal Facility under SPF conditions. All procedures were approved by New York University School of Medicine Institutional Animal Care and Use Committee.

### Adipose tissue stromal vascular fraction (SVF) purification

To isolate leukocytes from adipose tissue mice were anesthetized with a mixture containing 12.5mg/ml ketamine, 2.5mg/ml xylazine, and 25 mg/ml acepromazine and intracardially perfused with PBS with 5 mM EDTA. After dissection, the fat tissue was minced and incubated for 35 min at 37°C under gentle agitation (150rpm) in cRPMI (10% FBS, 1mM of Sodium Pyruvate (Gibco, 11360-070), Glutamax 1x (Gibco, 35050-061), MEM non-essential amino acids (Gibco, 11140-050), Hepes (Gibco, 15630-080), Pen Strep (Gibco, 15140-122) and RPMI (Gibco, 21870-076) containing 1mg/ml of collagenase type VIII (Sigma-Aldrich, C2139) and 100ug/ml of DNase I (Sigma-Aldrich, 10104159001). After digestion, the suspension was washed with ice-cold cRPMI and spun down. The floating fat layer was removed and the leukocyte-enriched pellet was ressuspended in cRPMI and filtered through a 70μm cell strainer.

### Isolation of mononuclear cells from spleen

Mice were perfused as described before. Spleens were minced and digested in PBS containing 5% of FBS, 10mM Hepes and 160 U/ml of collagenase D (Sigma-Aldrich, 11088882001) at 37°C for 35 min. After incubation 10 mM EDTA were added for 5 min in to disrupt DC–T cell complexes. The cell suspension was then filtered through a 70μM cell strainer.

### Bacteria

*Salmonella enterica* serovar Typhimurium (SL1344) strain (Yu et al., 2014) was used in infections experiments. Mice were injected i.v. with 10^8^ CFU and analyzed 3-6 hours post infection, as indicated. Briefly, a single aliquot of *Salmonella* was grown in 3 mL of LB overnight at 37°C with agitation, and then the bacteria were sub-cultured (1:30) into 3 mL of LB for 3h at 37°C with agitation. Bacteria were injected i.v. into recipient mice in a total volume of 100μl. *Staphylococcus aureus* strain Newman were cultured overnight from a single colony in 5 ml Tryptic Soy Broth (TSB) with shaking at 37°C. Cultures were diluted 1:100 in TSB (5 ml for i.v. infection for infection) and cultured shaking at 37°C for an additional 3h until optical density at 600 nm was 1.5 (10^9^ CFU/ml). Intravenous infection was performed via retro-orbital vein. Mice were injected i.v. with 10^8^ CFU and analyzed 12 hours post infection, as indicated.

### Antibodies and fluorescent conjugates

Antibodies were purchased from BD-biosciences (anti-active caspase 3 (clone C92-605) BUV395, 564095; anti-BrdU (clone 3D4) Percp-cy5-5, 560809; anti-CD45 (clone 30-F11) BUV 395, 564279; anti-Ly6G (clone 1A8) Percp-cy5.5, 560602; anti-SiglecF (clone E50-2440) BV421 and APC-R700, 565934 and 565183; Streptavidin-BUV395, 564176), Thermofisher (anti-CD8 (clone 53-6.7) APC-E780, 47-0081-82; anti-CD11b (clone M1/70) APC-E780, 47-0112-82; anti-IL33R (clone RMST2-2) PE, 12-9335-82; anti KI-67 (clone SolA15) PE, 12-5698-82; Ovalbumin-A647, O34784; Isolectin GS-IB4 A647, I32450; Qtracker 655 vascular labels, Q21021MP; Hoechst 33342, 62249. 4’,6-diamidino-2-phenylindole (DAPI), D3571 or Live/dead fixable blue, L34961 were used to exclude dead cells.), Biolegend (anti-CD4 Percp-cy5.5 (clone RM4-5), 100540; anti-CD11b (clone M1/70) A488 and A647, 101217 and 101218; anti-CD11c (clone N418) A647, 117312; anti-CD19 (clone 1D3) FITC, 152404; anti-CD31 (clone 390) A647, 102415; anti-CD45 (clone 30-F11) A700, 103128; anti-CD64 (clone x54-5/7.1) APC, PE, BV421 and Biotin, 139306, 139304, 139309 and 139318; anti-CD206 (clone C068C2) A647, PE-CY7 and PE/DAZZLE594, 141711, 141720 and 141731; anti-CD301 (clone LOM-14) Percp-cy5-5, 145709; anti-F4/80 (clone BM8) BV421, 123137; anti-I-A/I-E (clone M5/114.15.2) A488, A647 and BV711, 107616, 107618 and 107643; anti-Ly6C (clone HK1.4) A488 and BV510, 128021 and 128033; anti-Ly6G (clone 1A8) PE/DAZZLE594, 127647; anti-NK1.1 (clone PK136) BV650 and PE/DAZZLE594, 108736 and 108747; anti-γδ TCR (clone GL3) FITC, 118105; anti-Thy1.2 (clone 30-H12) FITC, 105306; anti-TCRβ (clone H57-597) BV510, 109233; Streptavidin-BV421; 405225) or Sigma-Aldrich (Rhodamine B isothiocyanate-dextran 70kDa; R9379)

### Flow cytometry and cell sorting

Flow cytometry data were acquired on an LSR-II flow cytometer (Becton Dickinson) and analyzed using FlowJo software (Tree Star 8.7). All stainings were performed as recommended by the antibody manufacturer. All samples were pre incubated with Fc block (2.4G2, BE0307 bioxcell) for 15 min at 4°C. FACS sorting was performed using an ARIA II sorter using a 100 μm nozzle (Beckton Dickinson). Flow cytometry of adipose tissue macrophages is made somewhat difficult because of their high autofluorescence. To minimize the impact of autofluorescence, we meticulously selected the fluorochromes for each cell marker, and used fluorescence minus one controls (FMOs).

### BrdU incorporation

WT mice were injected i.p. with 1 mg of BrdU twice, separated by 12 hours. Spleens and fat pads were harvested and processed as described before. Cells were surface-stained, and intranuclearly stained for BrdU using the BD BrdU proliferation kit (BD 552598).

### Monocyte transfer into HFD treated animals

To evaluate if adipose tissue macrophages subpopulations have origin from circulating monocytes on High Fat Diet-treated animals, monocytes were isolated from the spleens and bone marrow of CD45.1 WT mice by flow cytometry sorting as CD45^+^CD90^-^CD19^-^NK1.1^-^ CD11b^+^LY6G^-^MHCII^-^Ly6C^HIGH^. Sorted cells (2 × 10^6^) were injected i.v. into CD45.2 WT mice. After 80 hours, adipose tissue cells were isolated and analysed by flow cytometry.

### RNA isolation and sequencing

Total RNA from sorted target cell populations was isolated using TRIzol LS (Invitrogen; 10296010) followed by DNase I (Qiagen, 79254) treatment and cleanup with RNeasy Plus Micro kit (Qiagen, 74034). RNA quality was assessed using pico bioanalyser chips (Agilent, 5067-1513). Only RNAs displaying a RNA integrity number (RIN) of 9 or higher were used for downstream procedures. RNAseq libraries were prepared using the Nugen Ovation Trio low imput RNA Library Systems V2 (Nugen; 0507-08) according to the manufacturer’s instructions by the NYU Genome Technology Center. Pooled libraries were sequenced as 50 nucleotide, paired-end reads on an Illumina HiSeq 2500 using v4 chemistry.

### RNAseq data quality assessment and visualization

Illumina sequencing adapters and reads with Phred quality scores lower than 20 were removed with Trimmomatic. Trimmed reads were mapped to the *Mus musculus* genome Ensembl annotation release 91 using STAR v2.5.3a with default settings. The number of reads uniquely mapping to each gene feature in the corresponding annotation file was determined using featureCounts. The resulting count tables were passed to R for further analyses.

### Differential expression and gene ontology analyses

Consistency between replicates was checked using principal component analysis (PCA) and Euclidean distance-based hierarchical clustering on normalized counts. Weakly expressed genes, defined as having less than 2 Fragment Per Kilobase of transcript per Million mapped reads (FPKM) in at least one group of replicates were removed from differential expression analysis. Counts normalization were performed using the method of Trimmed Mean of M-values (TMM) from edgeR package. Differential expression analyses were performed using limma. Genes were considered significant when the observed log2 fold-change was greater than 1 and the adjusted P value was less than 0.05. The set of significant genes for each contrast of interest was passed to Goseq for functional gene categories enrichment analyses.

### Cell morphology

Sorted cells were spun onto Shandon Cytoslides with a Thermo Shandon Cytospin 4 cytofuge (10 min at 500 rpm), fixed with methanol and stained with Wright-Giemsa (Sigma-Aldrich; GS500). Images were acquired in a Zeiss AxioObserver microscope using a magnification of 63X N.A. 1.4 in bright field at room temperature. Zen software was used for image acquisition and ImageJ software, Fiji version 1.0, for contrast, brightness and pseudo-color adjustments.

### Confocal microscopy of WAT macrophages

White adipose tissue was stained as described (Martinez-Santibanez et al., 2014), with modifications. Mice were euthanized and slowly perfused by intracardiac injection with 20 ml of PBS/5mM EDTA. Epididymal fat pads were excised and fixed for 1h in PBS/1% paraformaldehyde with gentle shaking at 4°C. After that, samples were blocked for 1 h in PBS/5% BSA (blocking buffer) with gentle rocking at room temperature. Primary antibodies were diluted in blocking buffer and added to fat samples for 18h at 4°C. Samples were then washed three times with cold PBS and incubated with blocking buffer-diluted Alexa 647-conjugated isolectin and appropriate antibodies with gentle rocking at room temperature. Nuclei were stained with DAPI for 5 min at room temperature. Fat pads were imaged at room temperature on an inverted confocal microscope (Zeiss 710 MP; optical lenses 25x N.A. 0.8 and 63x N.A. 1.4) by placing the pad in Fluoromount-G (Southern Biotech, 0100-01) in a chambered coverslip. Zen software was used for image acquisition and ImageJ software, Fiji version 1.0, for contrast, brightness and pseudo-color adjustments.

### Adipo-Clear

Sample Collection: the method was performed as described before (Chi et al., 2018). Mice were injected with 18mg/kg of dextran-tetramethylrhodamine lysine fixable (Thermofisher, D1818). After one hour, animals were anesthetized as described before and an intracardiac perfusion/fixation was performed with 1xPBS followed by 4% PFA. All harvested samples were post-fixed in 4% PFA at 4°C overnight. Fixed samples were washed in PBS for 1 hr three times at RT.

Delipidation and Permeabilization: fixed samples were washed in 20%, 40%, 60%, 80% methanol in H2O/0.1% Triton X-100/0.3 M glycine (B1N buffer, pH 7), and 100% methanol for 30 min each. Sample were then delipidated with 100% dichloromethane (DCM; Sigma-Aldrich) for 30 min three times. After delipidation, samples were washed in 100% methanol for 30 min twice, then in 80%, 60%, 40%, 20% methanol in B1N buffer for 30 min each step. All procedures above were carried out at 4°C with shaking. Samples were then washed in B1N for 30 min twice followed by PBS/0.1% Triton X-100/0.05% Tween 20/2 μg/ml heparin (PTxwH buffer) for 1hr twice before further staining procedures.

Immunolabeling: samples were incubated in primary antibody dilutions in PTxwH for 4 days. After primary antibody incubation, samples were washed in PTxwH for 5 min, 10 min, 15 min, 30 min, 1 hr, 2 hr, 4 hr, and overnight, and then incubated in secondary antibody dilutions in PTxwH for 4 days. Samples were finally washed in PTwH for 5 min, 10 min, 15 min, 30 min, 1 hr, 2 hr, 4 hr, and overnight.

Tissue Clearing: samples were embedded in 1% agarose in PBS, and dehydrated in 25%, 50%, 75%, 100%, 100% methanol/H2O series for 30 min at each step at RT. Following dehydration, samples were washed with 100% DCM for 30 min twice, followed by an overnight clearing step in dibenzyl ether (DBE; Sigma-Aldrich). Samples were stored at RT in the dark until imaging. 3D Imaging: all whole-tissue samples were imaged on a light-sheet microscope (Ultramicroscope II, LaVision Biotec) equipped with 1.3X (used for whole-tissue views with low-magnification) and 4X objective lenses (used for high-magnification views) and an sCMOs camera (Andor Neo). Images were acquired with the ImspectorPro software (LaVision BioTec). Samples were placed in an imaging reservoir filled with DBE and illuminated from the side by the laser light sheet. The samples were scanned with the 488 nm, 640 nm and 790 nm laser channels and with a step-size of 3 μm for 1.3x objective and 2.5 μm for 4x objective.

Image Processing: all whole-tissue images were generated using Imaris x64 software (version 8.0.1, Bitplane). 3D reconstruction was performed using the “volume rendering” function. Optical slices were obtained using the “orthoslicer” tool. Optical sections were generated using the “snapshot” tool.

### Multiphoton microscopy

WT mice were anesthetized with i.p. injection of 20 μl/g of 2.5% Avertin before surgery for intravital imaging. Anesthesia was maintained by continuous administration 1% of isofluorane and 1l/min oxygen mixture while imaging was performed. Mice were injected with Hoechst dye (blue) and Qtracker 655 vascular labels for visualization of cell nuclei and blood vessels. qDots have a PEG surface coating especially designed to minimize interactions, and do not get taken up by VAMs or any other cell type in the adipose tissue. 10 min following induction of anesthesia, mice were placed on a custom platform heated to 37°C. Upon loss of recoil to paw compression, a small incision was made in the abdomen. One perigonadal fat pad was located, exposed and placed onto a raised block of thermal paste covered with a wetted wipe. A coverslip was placed on top of the fat. The platform was then transferred to the FV1000MPE Twin upright multiphoton system (Olympus) heated stage. Time-lapse was ± 30sec with a total acquisition time between 15 min to 1 hour depending of the condition. A complete Z stack (80 μm) was made during each acquisition. Depending on the experimental condition 450μg of dextran-rhodamine 70kDA was injected i.v. after 15 min of video acquisition.

### Computational analysis of multiphoton microscopy

For each individual movie, time-lapse was ± 30sec with a total acquisition time varying from 15min to 1 hour. Raw data as imported from the microscope was used for all subsequent analyses. *Imaris* (Bitplane AG) software was used for preparation of individual images and movies. Some post-analysis pseudo-color adjustment was performed for individual images and movies to account for differences in auto-fluorescence and Hoechst labeling. For all comparison images and movies (comparing pseudo-color channels showing dextran-rhodamine labeling over time) identical settings for all image parameters were used.

### Statistical analysis

Mean, SD and SEM values were calculated with GraphPad Prism (GraphPad Software). Error bars represent ± SEM. Unpaired Student’s *t* test was used to compare two variables. For 3 or more variables one way anova was used with a turkey post test, as indicated in each figure legend. P-values < 0.05 were considered significant. Statistics symbols were determined as: * = p<0.05, ** = p<0.01 and *** = p<0.001.

### Online Supplementary Materials

We have eight supplemental figures, one supplemental table, and three movies.

Fig S1: The epididymal white adipose tissue (eWAT) contains a broad representation of innate and adaptive immune cells.

Fig S2: Vascular-associated Adipose tissue VAM macrophages constitute the main immune cell population in the WAT

Fig S3: VAMs from mice fed normal diet display lipid droplets in their cytoplasm.

Fig S4: eWAT monocytes rapidly adapt their gene expression to the adipose tissue environment.

Fig S5: VAMs present antigens captured from blood circulation to T cells.

Fig S6: eWAT macrophages from HFD-fed mice do not upregulate expression of genes involved in the hypoxia response.

Fig S7: Systemic Gram-positive bacterial infection induces rapid changes in eWAT macrophages that are similar to those caused by Gram-negative bacteria.

Fig S8: β3 adrenergic receptor agonist induces remarkable adipose tissue neutrophilia.

Table S1: List of official and unofficial gene names and aliases.

Movie1: Close association between VAMs and adipose tissue blood vessels. Movie2: Rapid uptake of blood-borne molecules by VAMs.

Movie3: Impaired uptake of rhodamine by macrophages from mice fed HFD.

## Acknowledgements

The authors declare no competing financial interests. We thank Drs. Carey Lumeng, Peter J. Murray, P’ng Locke and Anthony W. Ferrante for helpful discussions at different stages of this project, and Maria Lafaille for valuable comments on the manuscript. We thank the kind assistance of Dr. Michael Cammer in helping with microscope acquisitions and image processing. We thank Rita Chan for preparing bacteria cultures for injection. We thank Dr. Gabriel Victora for kindly allowing the use of the FV1000MPE Twin upright multiphoton system (Olympus). We thank Dr. Dan Littman for generously sharing equipment, and critical review of the manuscript. The NYU microscope core is supported by grant NCRR S10 RR023704-01A1. NYU genomic core is a shared resource partially supported by the Cancer Center Support Grant, P30CA016087, at the Laura and Isaac Perlmutter Cancer Center. H.M.S. was supported by a fellowship from the National Council for Scientific and Technological Development (CNPq) (Brazil), and a postdoctoral fellowship from Dr. Bernard B. Levine.

## Supplemental Figure Legends

**Supplemental Figure 1. The epididymal white adipose tissue (eWAT) contains a broad representation of innate and adaptive immune cells.** Relevant to main Fig. 1**. (A-L)** Flow cytometry labeling of eWAT from 12-20 w.o. WT C57BL/6J male mice, showing the gating strategies. The thicker line gates of Fig. S1 H-L are color-coded, identifying with the same color each population throughout the manuscript. **(M)** Surface protein expression of several known myeloid cell markers. The histograms show a stack of 8 populations, identified as follows. From top to bottom: **CD11b^-^DCs**: Dendritic Cells CD45^+^CD90^-^CD19^-^CD11b^-^MHCII^+^CD11c^+^. **Eos**: Eosinophils CD45^+^CD90^-^CD19^-^CD11b^+^CD206^LOW/INT^SiglecF^+^. **Neu**: Neutrophils CD45^+^CD90^-^CD19^-^ CD11b^+^CD206^LOW/INT^Ly6G^+^. **CD11b^+^DCs**: Dendritic cells CD45^+^CD90^-^CD19^-^CD11b^+^CD206^LOW/INT^ MHCII^+^CD64^-^CD11c^+^. **eMon**: eWAT Monocytes CD45^+^CD90^-^CD19^-^CD11b^+^CD206^LOW/INT^SiglecF^-^Ly6G^-^MHCII^NEG/LOW^Ly6C^HIGH^. **DPs**: Double positive macrophages CD45^+^CD90^-^CD19^-^CD11b^+^CD206^LOW/INT^MHCII^+^CD64^+^CD11c^+^. **PreVAM**: Pre Vasculature-associated Adipose tissue Macrophages CD45^+^CD90^-^CD19^-^CD11b^+^CD206^LOW/INT^MHCII^+^CD64^+^CD11c^-^. **VAM1**: Vasculature-associated Adipose tissue Macrophages subtype 1 CD45^+^CD90^-^CD19^-^ CD11b^+^CD206^HIGH^MHCII^HIGH^Tim4^INT^. **VAM2**: Vasculature-associated Adipose tissue Macrophages type 2 CD45^+^CD90^-^CD19^-^CD11b^+^CD206^HIGH^MHCII^INT^Tim4^HIGH^. SVF: Stromal vascular fraction. FSC: forward scatter. SSC: Side scatter. Displayed above each dot plot is the sub-gate in which the population is found. Numbers inside the plots represent percentages. Representative data of at least 2 independent experiments with 3 animals on each.

**Supplemental Figure 2. Vascular-associated Adipose tissue VAM macrophages constitute the main immune cell population in the WAT.** Relevant to main Fig. 1**. (A)** Distribution of all immune cell types per gram of eWAT from WT mice age 12-20 w.o. fed normal diet. **(B)** Number and percentage of macrophages, dendritic cells and monocytes in three different white adipose tissues depots. **mWAT:** Mesenteric white adipose tissue. **eWAT:** Epididymal white adipose tissue. **sWAT:** Subcutaneous white adipose tissue. SVF: Stromal vascular fraction. *: p≤0.05. **: p≤0.01. ***: p≤0.001. Statistical analysis was carried out using one way anova followed by a turkey post test. Data displayed is the compilation of at least 3 independent experiments with 3 animals on each.

**Supplemental Figure 3. VAMs from mice fed normal diet display lipid droplets in their cytoplasm.** Relevant to main Fig. 1. eWAT myeloid populations were sorted by flow cytometry, prepared on glass slides by Cytospin, stained with Wright-Giemsa, and visualized in a microscope set to bright field. 63x magnification. **(A-E)** Morphology of VAMs, eMon, PreVAMs, DP macrophages, eosinophils, CD11b^+^DC and CD11b^-^DC, as described in **Fig.S1A.** bDC abbreviates CD11b^+^ dendritic cells.

**Supplemental Figure 4. eWAT monocytes rapidly adapt their gene expression to the adipose tissue environment.** Relevant to main Fig. 1**. (A)** Comparative analysis of eWAT gene expression compared to blood Ly6C^+^ and Ly6C^-^ monocytes. eMon RNAseq data was compared to publically available RNAseq data for blood Ly6C^+^ and Ly6C^-^ monocytes (Mildner et al., 2017). Each column represents consolidated data from 3 biological replicates per cell type. The z-score of the gene expression profiles gives a scale to measure the differential expression. **(B)** Heatmap displaying differentially expressed genes between eWAT monocytes and Ly6C^+^ and Ly6C^-^ blood monocytes. **(C)** Expression of typical dendritic cell (DC) genes in the eWAT myeloid cells. **B and C** display gene expression (FPKM in the Log_10_ base). Yellow box in Fig. S4C marks the two dendritic cell populations.

**Supplemental Figure 5. VAMs present antigens captured from blood circulation to T cells.** Relevant to main Fig. 2. Sterile whole ovalbumin-A647 (Ova) was injected i.v. in WT mice. Twelve hours post injection, VAMs were purified by flow cytometry and cocultured with Ova-specific OTII T cells. Twelve hours later, T cell activation was determined by the upregulation of CD69 expression.

**Supplemental Figure 6. eWAT macrophages from HFD-fed mice do not upregulate expression of genes involved in the hypoxia response.** Relevant to main Fig. 3**. (A)** RNAseq analysis of genes associated with the hypoxia response of the sorted myeloid populations obtained from WT mice under normal diet (ND) or after 16 weeks of HFD feeding. Data are displayed as gene expression in FPKM (Log_10_ base). **(B)** RNAseq analysis of genes associated with M1, M2, or MMe of the sorted myeloid populations obtained from WT mice under normal diet (ND) or after 16 weeks of HFD feeding. Data are displayed as gene expression in FPKM (Log_10_ base).

**Supplemental Figure 7. Systemic Gram-positive bacterial infection induces rapid changes in eWAT macrophages that are similar to those caused by Gram-negative bacteria.** Relevant to main Fig. 5**. (A-B)** Analysis of the spleen of LPS-injected mice. WT mice were injected i.v. with 50μg of LPS and splenic myeloid cells distribution was determined by flow cytometry 24h post injection. **(A)** representative dot plot **(B)** percentages of the indicated myeloid cell populations. Representative data of 3 independent experiments (*n*= 3-5 mice). Eos: Eosinophils. Neu: Neutrophils. DC: Dendritic cell. Mon: Monocytes. MZ Mac: Marginal zone macrophages. **(C)** Active caspase 3 positive cells from the experiment shown in main **Fig. 5C. (D-E)** Analysis of the spleen of *Salmonella*-infected mice. Same animals as shown in main **Fig. 5H-J. (D)** representative dot plot **(E)** percentages of the indicated myeloid cell populations. **(F-J)** Analysis of *Staphylococcus aureus* (S.aureus)-infected mice. WT mice were injected i.v. with 10^8^ CFU. of S.aureus and sacrificed 12h post infection for eWAT and spleen analysis. **(G-J)** Flow cytometry analysis of splenic and eWAT myeloid cell populations. **(G)** Representative eWAT dot plots showing macrophage populations; **(H)** Absolute number of myeloid cell populations; **(I)** Representative dot plots showing splenic myeloid cells; **(J)** Percentage of splenic myeloid cells; Representative data of 2 independent experiments (*n*= 3-5 mice). *: p≤0.05. **: p≤0.01. ***: p≤0.001. Statistical analysis was carried out using one way anova together with turkey post test or unpaired T test (only for pair comparisons). AT: adipose tissue.

**Supplemental Figure 8. β3 adrenergic receptor agonist induces remarkable adipose tissue neutrophilia.** Relevant to main Fig. 6**. (A-D)** Animals were fasted for one day, in cages with paper-based bedding. **(A)** Kinetics of serum non-esterified fatty acid (FFA) content during 24h fasting (*n*=5). **(B-C)** Blood FFA and glucose levels of the animals analyzed in main **Fig. 6A-B. (D)** Daily weight variation of mice shown in main **Fig. 6C-D**. **(E)** Flow cytometry analysis of active caspase 3-positive cells of the experiment shown in main **Fig. 6A-B. (F-H)** Representative dot plots from the experiment shown in main **Fig. 6G-H** showing the accumulation of **(G)** neutrophils (Ly6G^+^) and **(H)** monocytes (Ly6C^+^MHCII^LOW^) in the eWAT of animals treated with the β3 adrenergic receptor agonist CL 316243.

## Movie Legends

**Movie 1. Close association between VAMs and adipose tissue blood vessels.** Light sheet microscopy of full epididymal fat pad post WAT clarification using the Adipo-Clear technique (Chi et al., 2018). CD31 (seen in red color) indicates blood vessels. Animals were injected with fixable dextran-rhodamine (tetramethylrhodamine, seen in green color), and, after one hour, animals were perfused to remove the dextran from the circulation, and submitted to the Adipo-Clear method. VAMs become labeled by their uptake of dextran-rhodamine. The movie shows the rotation of a full epididymal fat pad. 4X maginification.

**Movie 2. Rapid uptake of blood-borne molecules by VAMs.** Intravital imaging of eWAT dextran-rhodamine uptake after i.v. injection. qDots (Qtracker 655) and Hoechst were i.v. injected immediately before the beginning of the recording, while dextran-rhodamine was injected while the recording was taking place, as indicated in the movie. qDots have a PEG surface coating especially designed to minimize interactions, and do not get taken up by VAMs or any other cell type in the adipose tissue. The arrow indicates a small field in which VAMs are uptaking dextran-rhodamine, an event that can be noticed as shortly as 1 minute after dye injection, and peaks in brightness at around 20 minutes. After dye injection, the imaging proceeds for 30 minutes, and it is looped once. 25x magnification.

**Movie 3. Impaired uptake of rhodamine by macrophages from mice fed HFD.** Intravital microscopy analysis of eWAT dextran-rhodamine uptake in mice fed ND or HFD. qDots (Qtracker 655), Hoechst, and dextran-rhodamine were i.v. injected and the uptake of rhodamine was observed. The first half of the movie shows a ND-fed mouse (1 hour of recording), followed by a HFD-fed mouse (also 1 hour). Not only the low rhodamine uptake can be observed, but also the overall disruption of normal tissue structure. 25x magnification.

